# Liver Fatty Acid Binding Protein (FABP1) Affects Cytochrome P450 2C9 (CYP2C9)-Mediated Metabolism in Human Liver Models

**DOI:** 10.64898/2026.10.12.758325

**Authors:** Yue Winnie Wen, Eli Friedman, Alex Zelter, King C. Yabut, Abhinav Nath, Michael J. MacCoss, Nina Isoherranen

## Abstract

Metabolic clearance of drugs is often systematically over- or underpredicted from human liver microsomes (HLM). We hypothesized that lack of consideration of interactions of cytosolic liver fatty acid binding protein (FABP1) with drugs and cytochrome P450s (CYPs) explains the off-prediction. We used two CYP2C9 probe substrates, diclofenac and tolbutamide, to test this hypothesis. We found that presence of FABP1 alters the kinetics of metabolite formation in HLM. The model that best fit the kinetic data included FABP1-CYP2C9 protein-protein interactions. Statistical modeling of proteomic expression and drug hydroxylation across a panel of 61 human liver S9 fractions (HLS9) showed that CYP2C9 and FABP1 expression together best predicted probe substrate hydroxylation. Incorporation of FABP1 expression and protein-protein interactions into novel physiologically based liver models significantly improved the quantitative accuracy of *in vivo* hepatic clearance prediction. Together, our data demonstrate the importance of taking FABP1 into consideration for CYP2C9-mediated metabolism.

## Introduction

Liver fatty acid binding protein (FABP1) is one of the most abundant cytosolic proteins expressed in the human liver.^1,2^ FABP1 constitutes 7–10% of total cytosolic protein.^1,2^ It is predominately expressed in the hepatocytes with some expression also found in the stellate cells.^3^ As a carrier of endogenous lipophilic ligands, FABP1 alters the metabolism and cellular distribution of fatty acids.^4–9^ Many drugs also bind to FABP1 with low micromolar equilibrium dissociation constants (K_d_s)^1,2,10–12^ and FABP1 has been suggested to alter drug distribution^10^ and drug metabolism^12–15^ in the liver. In particular, FABP1 appears to alter the efficiency of CYP mediated metabolism when studied with purified or recombinant CYP enzymes.^12,13,16^ Presence of FABP1 in incubations in comparison to absence of FABP1 resulted in decreased intrinsic clearance of diclofenac by recombinant CYP2C9^16^ and of (-)-trans-Δ⁹-tetrahydrocannabinol (THC) by recombinant CYP2C9, CYP2C19, and CYP3A4.^14^ FABP1 was shown in a separate study to interact with purified CYP3A4 with nanomolar affinity, and kinetic modeling of the data favored a model of directed substrate transfer from FABP1 to CYP3A4.^12^ However, quantitative translation of these protein-drug and protein-protein interactions of FABP1 to *in vivo* drug clearance prediction has not been defined, and the presence of these interactions in more complex models such as human liver microsomes (HLM) or human liver S9 fractions has not been shown.

A highly prevalent T to C single nucleotide polymorphism (SNP) in exon 3 of the human FABP1 gene results in a threonine-to-alanine amino acid change at position 94 (p.Thr94Ala; T94A) of the protein.^17–19^ The minor allele frequency of the C allele coding for the A94 protein is 26–38% in the human population^17–20^. The substitution of the polar, β-branched threonine by a smaller and nonpolar alanine in the β-strand G in FABP1, occupies approximately 24% less volume.^21^ Although the threonine residue at position 94 is highly conserved across mammalian FABP1 including humans, rats, mice, pigs, and cattle,^19^ the alanine substitution does not appear to affect binding affinity for fatty acids, triglyceride synthesis intermediates, or fibrates.^17,21^ Instead, A94 SNP has been suggested to result in changes in conformational response of FABP1 to ligand binding, subsequently impairing nuclear entry and peroxisome proliferator-activated receptor α (PPARα) activation. These functional changes with A94 FABP1 may explain the dyslipidemia, metabolic dysfunction-associated fatty liver disease (MAFLD), and fenofibrate non-response associated with the A94 Proteoform.^17–20^ Further the alanine substitution in human FABP1 may affect protein-protein interactions of FABP1 in the cytosol. Previous studies of FABP1-CYP interactions and the impact of FABP1 on liver distribution of drugs have only considered the wild-type (WT) proteoform.^10,12–14,16^ Hence, there is a need to define whether T94A substitution affects drug disposition and the interpretation of FABP1 effects on drug distribution and metabolism.

*In vitro* metabolic data from human liver preparations often systematically underpredicts *in vivo* hepatic clearance,^22,23^ suggesting some disposition mechanisms are missing in the *in vitro* experiments.^24^ Nonspecific binding to microsomal lipids, loss of cytosolic proteins and physiological macromolecular crowding, or unstirred water layer and permeability limitations in hepatocyte suspensions have been proposed as potential reasons for the underprediction.^25–27^ The data collected from subcellular fractions containing HLM and whole cell hepatocyte experiments were reported to disagree for intrinsic clearance predictions for CYP3A4 substrates.^28^ For CYP2C9 substrates, HLM data systemically underpredict *in vivo* clearance.^29–31^ We hypothesized that the lack of FABP1 in the *in vitro* HLM system and the lack of incorporation of distribution and substrate channeling effects of FABP1 in HLM play a role in the underprediction of hepatic clearance of CYP2C9 substrates. We chose diclofenac and tolbutamide, two CYP2C9 probe drugs that bind to FABP1 but have minimal microsomal binding, as model drugs to test this hypothesis. The goal of this study was to determine, using relevant human liver model systems, whether WT or A94 substituted FABP1 alter substrate metabolism and liver distribution and to test using a novel physiologically based liver model whether incorporation of FABP1 into the model allows for more accurate *in vitro* to *in vivo* clearance predictions.

## Results

### FABP1 WT and A94 display similar ligand-binding behavior

The binding of the fluorescent fatty acid probe of FABP, 11-(dansylamino)undecanoic acid (DAUDA), to WT and A94 FABP1 was characterized (**Supplementary Fig. 1)**. The best-fit binding affinities of DAUDA (K_d,1_ = 194 ± 41 nM for WT; K_d,1_ = 185 ± 13 nM for A94) were not different between the WT and A94 FABP1 when FABP1 was titrated into a solution of DAUDA (**Supplementary Fig. 1)**. When DAUDA was titrated to FABP1, the K_d,2_ was >10 µM for both proteoforms. The DAUDA binding kinetics were further characterized using stopped-flow fluorescence (**Supplementary Fig. 2; Supplementary Table 1**). Similar to the fluorescence titrations, the binding constants of DAUDA to WT and A94 FABP1 were comparable (**Supplementary Table 1**). The kinetic K_d_ values agreed well with the equilibrium K_d,1_ from fluorescence titration for DAUDA. Together, these data show that the T94A substitution in FABP1 does not affect the rate of the fluorescent probe association nor the dissociation from FABP1.

The binding of several drugs to WT and A94 variant FABP1 was similar when measured by fluorescence displacement assays with DAUDA as a probe (**Fig. 1** and **Table 1**). None of the drug ligands fully abolished the DAUDA–FABP1 fluorescence (**Fig. 1** and **Supplementary Fig. 3**). Instead, these ligands attenuated the probe fluorescence partially, and the residual spectra remained blue-shifted relative to free probe (**Fig. 1** and **Supplementary Fig. 3**). This suggests formation of a ternary ligand–FABP1–DAUDA complex as previously described and characterized.^16^ The K_d_ values for the drug ligands were in the low micromolar range with no significant differences observed in K_d_ values between WT and A94 FABP1, indicating minimal effects of the T94A substitution on the binding affinities for the drugs tested (**Table 1**).

**Figure 1.**
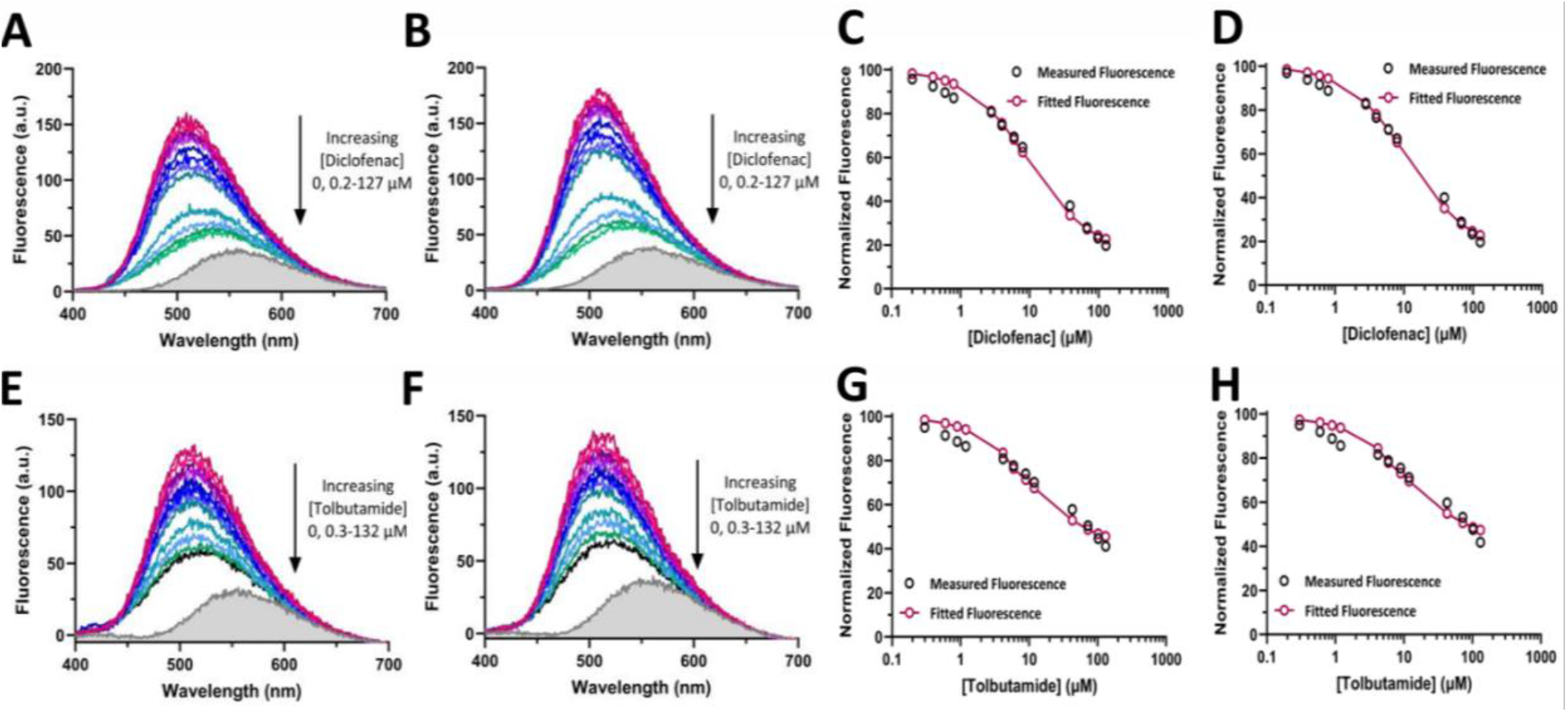
DAUDA displacement from WT or A94 FABP1 by diclofenac and tolbutamide. **A-B** The raw fluorescence spectra for DAUDA displacement titrations with WT protein **(A)** or A94 FABP1 **(B)** with diclofenac as ligand. **C-D** The corresponding DAUDA displacement curves with WT FABP1 **(C)** or A94 FABP1 **(D)** for panels **A-B**. **E-F** The raw fluorescence spectra for DAUDA displacement titrations with WT protein **(E)** or A94 FABP1 **(F)** and tolbutamide as ligand. **G-H** The corresponding DAUDA displacement curves with WT FABP1 **(G)** or A94 FABP1 **(H)** for panels **E-F**. The concentration ranges of drug ligands titrated are indicated next to the raw fluorescence spectra. The gray shaded areas show the spectrum of 0.5 μM unbound DAUDA in the absence of FABP1 and drug ligand. Solid lines on the panels on the right indicate fits to a ternary complex binding model as previously established.^16^ The resulting K_d_ values are summarized in Table 1.

**Table 1.** Summary of the FABP1 binding affinities of select drugs measured by DAUDA displacement. All values are reported as means ± standard deviation from three replicate experiments conducted on separate days. The p values were estimated using paired t-tests.

| Drug | WT $K_d$ (μM) | A94 $K_d$ (μM) | P value |
| --- | --- | --- | --- |
| Diclofenac | $5.2 \pm 1.3$ | $7.2 \pm 1.2$ | 0.18 |
| Gemfibrozil | $8.6 \pm 0.9$ | $10.9 \pm 0.7$ | 0.13 |
| Pioglitazone | $2.4 \pm 0.4$ | $2.9 \pm 0.6$ | 0.11 |
| Tolbutamide | $7.4 \pm 2.3$ | $7.2 \pm 2.0$ | 0.94 |

### FABP1 WT and A94 display similar conformational stability

To compare the conformational stability of the two proteoforms, thermal denaturation was monitored by Far-UV circular dichroism (CD) across a temperature ramp (35–95 °C). A two-state unfolding model^32^ was fit to the normalized fraction unfolded data at 222 nm^32,33^ wavelength (**Supplementary Fig. 4**). Both WT and the A94 variant FABP1 unfolded through a single cooperative transition with nearly identical melting temperatures (T_m_ = 69.5 ± 3.9 °C for WT and 69.3 ± 4.6 °C for A94). The unfolding curves appeared superimposable across the full temperature range. Thus, FABP1 A94 likely retains the thermal stability of the WT protein.

### Characteristics of human liver bank and FABP1 expression across different FABP1 genotypes and liver pathologies

Human liver samples from individual donors were obtained from the University of Washington (UW) human liver bank.^34,35^ A total of 61 human livers were included in the study. Demographic information of these human liver donors and details of liver pathology can be found in **Supplementary Table 2**. To test whether the FABP1 promoter SNP (rs2919872 chr2:88129052 C>T) or the FABP1 A94 coding SNP (rs2241883 chr2:88124547 T>C) correspond to altered FABP1 expression, the 61 human livers included in the study were genotyped for the two SNPs. The FABP1 promoter C allele was associated with the wild type T allele of FABP1 based on the chi-square test of independence (p = 0.00018, Cramér’s V = 0.427, **Supplementary Fig. 5**). The total FABP1 expression was previously quantified and reported^10^ by us using the same set of 61 human liver S9 fractions (HLS9). The FABP1 content from those data ranged from 0.60 to 1.64 nmol/mg S9 protein.^10^ These protein expression data allowed us to evaluate the impact of the FABP1 SNPs on total FABP1 expression in the liver bank. FABP1 expression was not different between the livers with different promoter genotypes (0.97 ± 0.23 nmol/mg S9 for C/C, 1.08 ± 0.26 nmol/mg S9 for C/T, and 0.98 ± 0.14 nmol/mg S9 for T/T, **Fig. 2A**). Similarly, FABP1 expression was comparable among the FABP1 T94A coding variant genotypes (0.98 ± 0.22 nmol/mg S9 for T/T homozygous, 1.06 ± 0.26 nmol/mg S9 for T/C heterozygous, and 0.97 ± 0.11 nmol/mg S9 for C/C homozygous, **Fig. 2B**).

**Figure 2.**
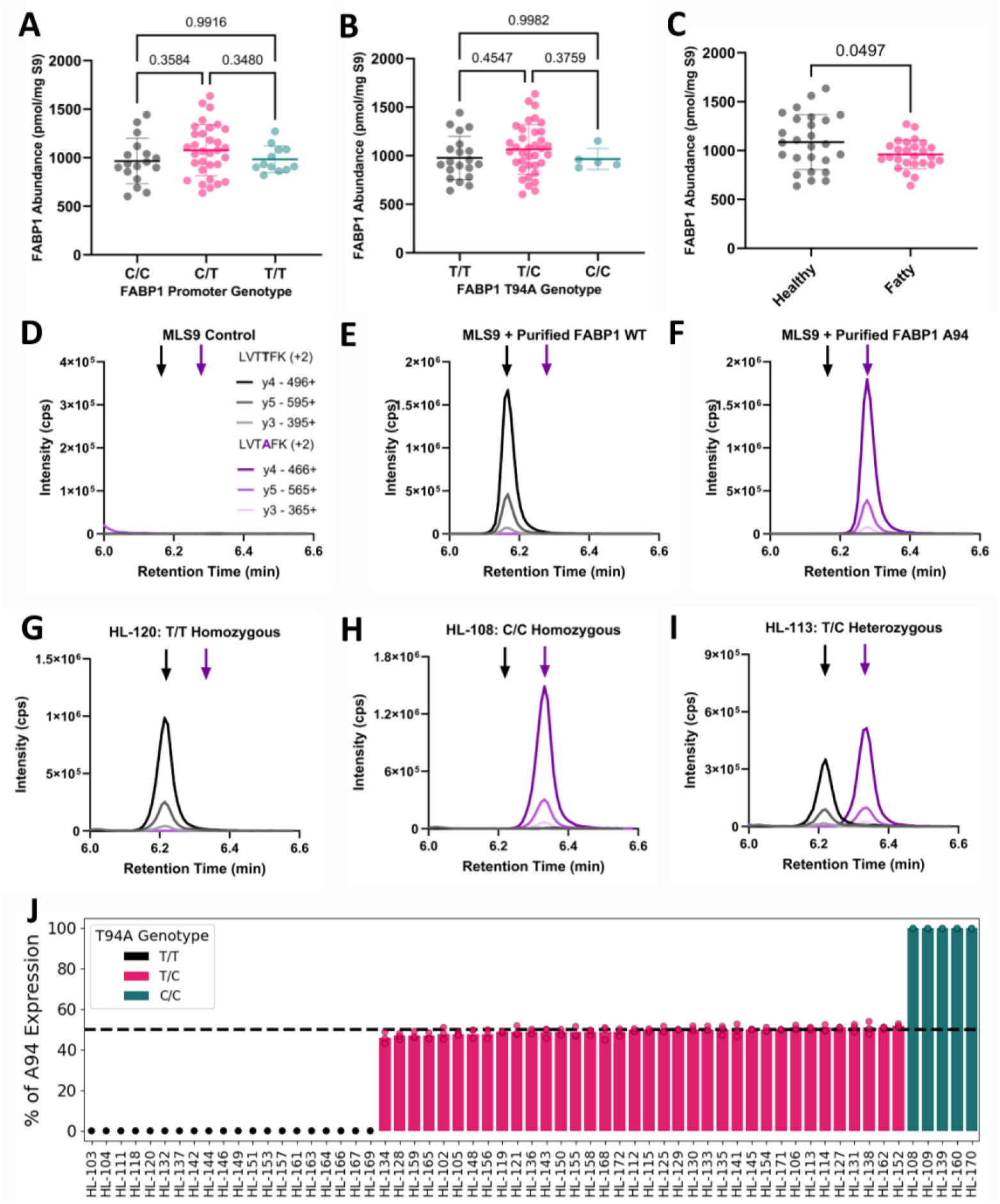
Total and proteoform level quantification of WT and A94 FABP1 expression in human livers across different FABP1 genotypes and with different liver pathology. Total FABP1 protein abundance (pmol/mg S9) separated by FABP1 promoter genotype **(A),** T94A genotype **(B)**, or liver pathology (normal versus fatty) **(C)** analyzed using donor records, previously reported^10^ total FABP1 expression data and genotyping data. Each point is an individual liver sample. Horizontal lines denote the mean ± standard deviation. Groups were compared by Kruskal-Wallis test with Dunn’s post hoc test and p values are shown above the brackets. Chromatograms of LVTTFK (black arrow) and LVTAFK (purple arrow) peptides from control MLS9 **(D)**, MLS9 spiked with 200 nM WT **(E)** or A94 **(F)** FABP1 show assay specificity. Representative chromatograms of FABP1 peptides LVTTFK and LVTAFK from T/T homozygous **(G),** T/C heterozygous **(H)**, and C/C homozygous **(I)** livers. The measured percentage of total FABP1 expressed as A94 variant in the 61 individual human livers analyzed is shown in **(J)** colored by genotype (pink T/C heterozygous, green C/C homozygous). The dashed line marks 50% of total FABP1 expression.

Some studies^18,19,36^ have reported association between FABP1 genotype and risk of metabolic dysfunction-associated steatotic liver disease (MASLD). Hence, we evaluated genotype enrichment in the fatty compared to healthy livers and compared FABP1 expression between healthy and fatty livers. Within our study of only 61 liver donors, A94 coding C allele carriers were more frequent among the fatty than normal livers, corresponding to an odds ratio of 2.08 (95% CI 0.62-6.97; Fisher’s exact p = 0.37). The FABP1 abundance in healthy livers (1.09 ± 0.28 nmol/mg S9) appeared higher (p = 0.0497) than in fatty livers (0.96 ± 0.15 nmol/mg S9) with the variance between the healthy and fatty livers being significantly different (p = 0.0018, **Fig. 2C**).

While the overall FABP1 expression in the human liver was not different amongst FABP1 genotypes, it is possible that different proteoforms represent different proportions of the liver FABP1 in heterozygous livers. To distinguish between these different FABP1 proteoforms, we developed a targeted LC-MS/MS based assay to quantify the relative abundance of the different FABP1 proteoforms, WT(T94) and A94, in the 61 human liver samples (**Fig. 2D-I**). This method allowed measurement of the concentrations of the tryptic surrogate peptides containing the polymorphic residue (LVTTFK for WT and LVTAFK for A94). Purified human T94 or polymorphic A94 FABP1 was spiked into mouse liver S9 fractions (MLS9) and digested as previously described.^10^ The detection of the specific peptides is shown in **Fig. 2D-F**. In digested HLS9 samples from livers with the TT genotype, only the LVTTFK peptide was detected. In livers genotyped as CC, only the LVTAFK peptide was detected (**Fig. 2G,H**). In heterozygous livers both peptides were detected (**Fig. 2I**). When the concentrations of LVTTFK and LVTAFK peptides were compared in the 36 FABP1 T94A heterozygous livers, the concentrations of LVTTFK and LVTAFK peptides were similar within the individual, indicating that in human liver there is no enrichment for either proteoform (**Fig. 2J**). The percentage of the A94 variant expression of the total FABP1 expression ranged from 46% to 51%, indicating equivalent protein transcription and stability for both FABP1 proteoforms *in vivo*.

### Simulation of liver tissue-to-plasma partition coefficient (K_p_) in the presence of WT and A94 FABP1

We have previously shown that FABP1 binding alters liver distribution of drugs that bind to FABP1.^10^ Expression levels of FABP1 WT and A94 in the 61 human liver samples were calculated from the total FABP1 expression^10^ and the percentage of A94 expression shown in **Fig. 2J**. Using the drug-FABP1 binding affinities determined in **Table 1**, the liver distribution of diclofenac, pioglitazone, gemfibrozil, and tolbutamide in the presence of the two FABP1 proteoforms was simulated (**Fig. 3**). When the expression of each proteoform and the binding affinity of the drugs to the WT and A94 FABP1 were incorporated into the model, the predicted liver K_p_ values were overall similar for the homozygous FABP1 WT livers and the homozygous A94 livers. The K_p_ values for diclofenac and pioglitazone trended lower in the presence of the A94 variant while gemfibrozil and tolbutamide were predicted to have similar K_p_ values (**Fig. 3A-D**).

**Figure 3.**
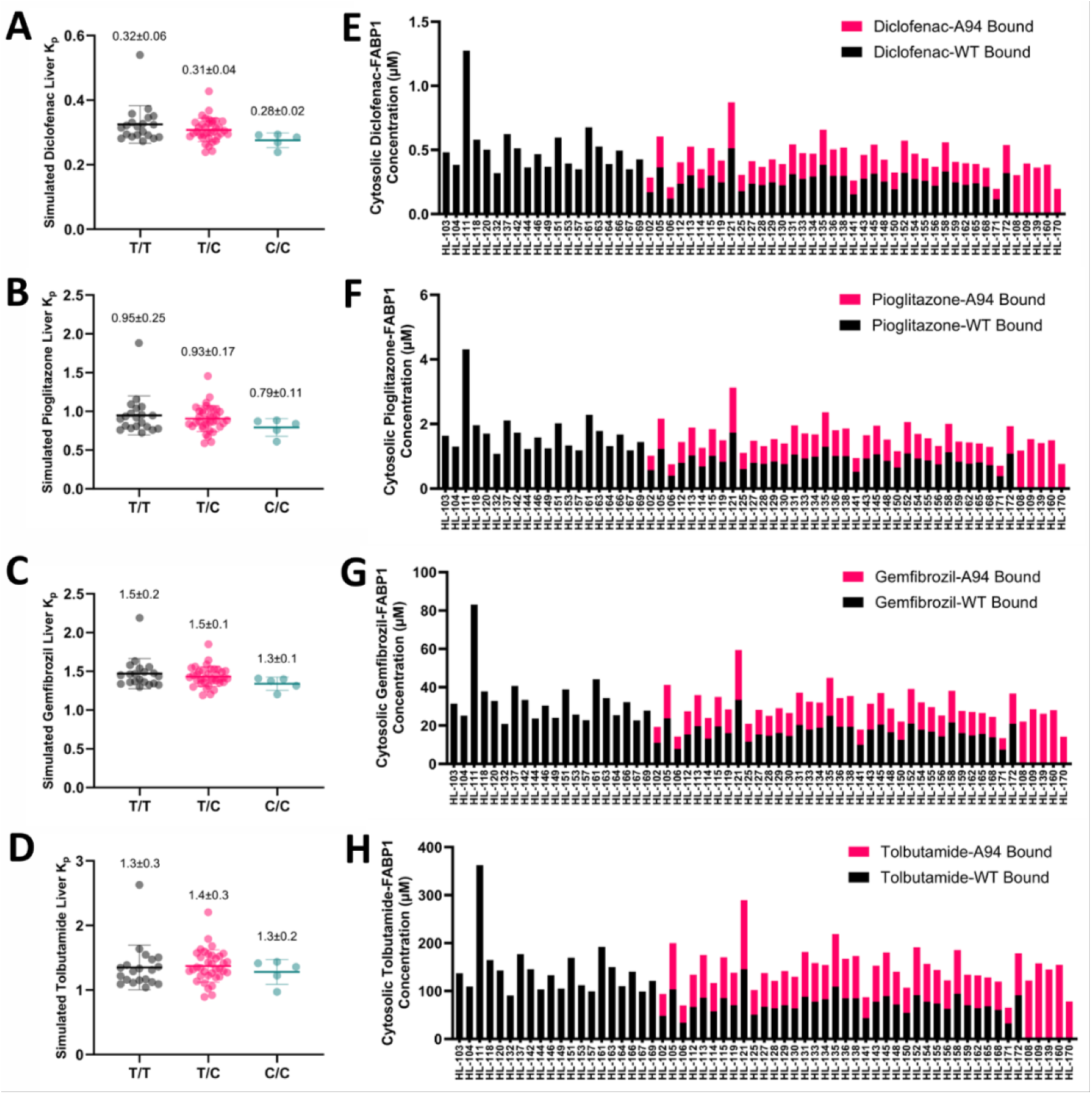
Predicted impact of FABP1 genotype on the extent of drug distribution to the liver. Simulated diclofenac **(A)**, pioglitazone **(B)**, gemfibrozil **(C)**, and tolbutamide **(D)** liver tissue-to-plasma partition coefficient (K_p_) separated by FABP1 T94A genotypes. The simulated K_p_ values for T/T (black circles), T/C (pink circles), and C/C (green circles) are shown for each liver with mean value listed above the data. The predicted cytosolic concentrations of WT and A94 FABP1-drug complex simulated based on measured K_d_ values and FABP1 expression levels in the 61 human livers for diclofenac **(E)**, pioglitazone **(F)**, gemfibrozil **(G)**, and tolbutamide **(H)**. The black bars show WT drug-FABP1 complex concentration and the pink bars show the cytosolic drug-FABP1 A94 bound concentrations.

Since the A94 proteoform has been proposed to interact differently with nuclear receptors than the WT FABP1, the relative abundance of cytosolic drug-FABP1 complex of each proteoform may have implications for downstream pharmacology. Hence, the intracellular, cytosolic, drug-FABP1 bound concentrations were simulated for the 61 human liver donors considering the individual donor genotype and proteoform expression (**Fig. 3E-H**). Overall, the coefficients of variation of intracellular drug-FABP1 bound concentrations were predicted to be between 33.0% to 35.6% for the four drugs (**Figure 3E-H**). The average unbound drug concentrations at steady state were predicted to be only 1.6%, 0.7%, 2.7%, and 3.1% of the FABP1-bound concentrations for diclofenac, pioglitazone, gemfibrozil, and tolbutamide respectively at clinically seen circulating concentrations as previously published.^10^

### FABP1 alters CYP2C9-mediated metabolite formation kinetics in pooled human liver microsomes (HLMs)

The WT FABP1 has been reported to affect CYP-mediated substrate metabolism with recombinant CYP enzymes,^12–14,16^ but these findings have not been translated to human liver fractions or to clearance predictions. To test whether FABP1 is important in modulating substrate metabolism, the impact of FABP1 WT and A94 variant on CYP2C9-mediated diclofenac and tolbutamide metabolism was evaluated. Diclofenac and tolbutamide were chosen as model drugs as they do not partition to lipid membranes that could confound data interpretation. In the absence of FABP1 they are completely unbound (f_u,mic_=1) in the HLMs.^26,37^.

Addition of 20 µM human FABP1 WT or A94 to pooled HLMs with varying concentrations of diclofenac or tolbutamide altered the formation kinetics of 4’OH-diclofenac (4’OH-DCF) and 4’OH-tolbutamide (4’OH-TBU) (**Fig. 4, Supplementary Fig. 6 and Supplementary Table 3**). In the presence of 20 µM FABP1, diclofenac binds to FABP1. Unbound concentrations of diclofenac in the presence of WT or A94 FABP1 were estimated using the K_d_ values determined in this study and a previously developed steady-state model taking each substrate and FABP1 concentration into account.^15^ The apparent unbound Michaelis-Menten constants (K_m,u_) that best fit the data between free diclofenac concentration and the product formation velocity were significantly lower when either FABP1 WT or A94 was present compared to no FABP1 controls (**Fig. 4B** and **Table 2**). However, maximum velocity (V_max_) of diclofenac hydroxylation was not affected by the presence of 20 µM FABP1 (**Fig. 4B** and **Table 2**). The unbound intrinsic clearance of diclofenac increased significantly (2.5- to 3.9-fold) in the presence of FABP1 WT or FABP1 A94 variant when compared to no FABP1 group. The unbound intrinsic clearance of diclofenac was also higher in the presence of WT FABP1 than A94 FABP1 likely due to the difference in diclofenac K_m,u_ between the FABP1 proteoforms.

**Figure 4.**
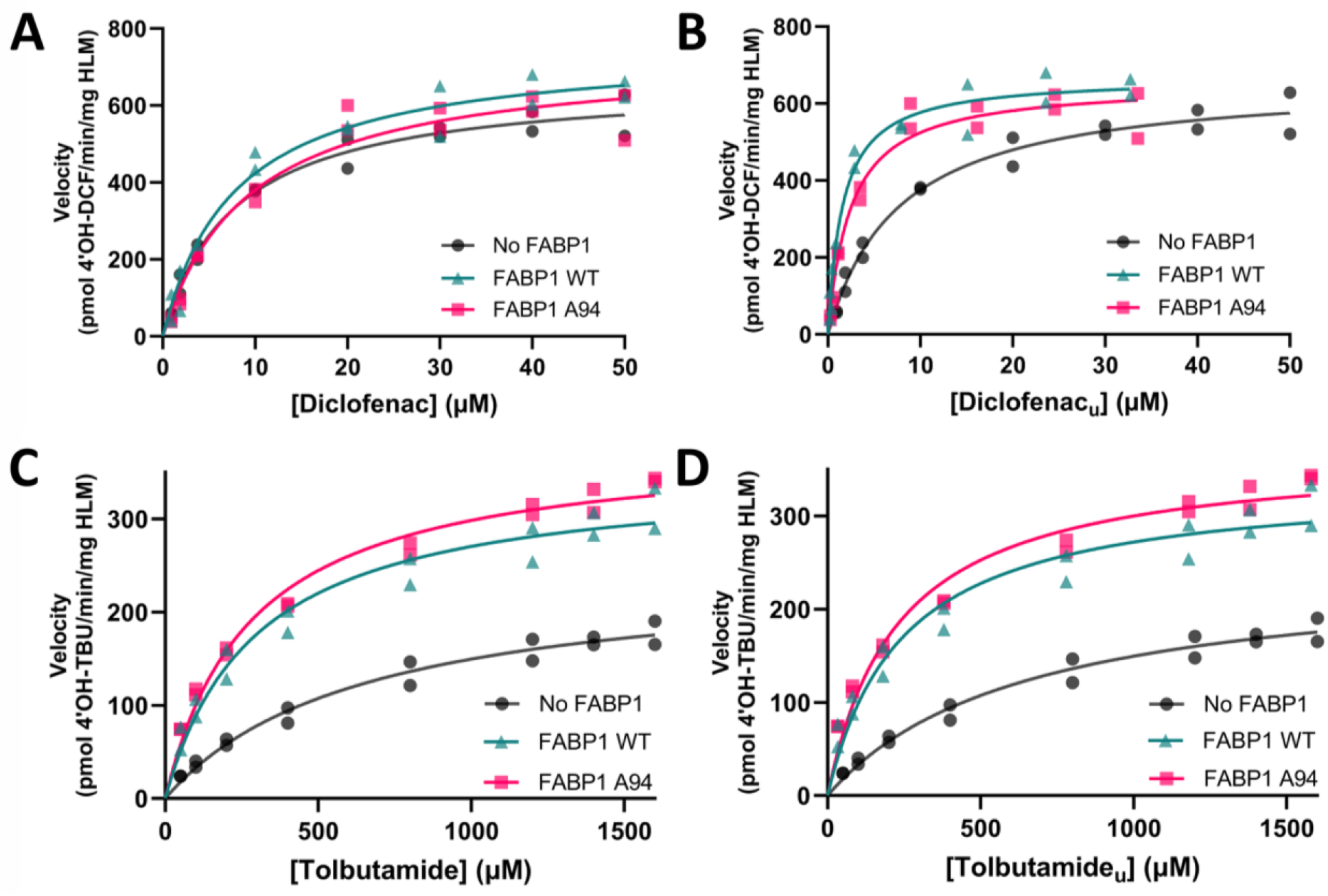
Effect of FABP1 on CYP2C9-mediated metabolism of diclofenac and tolbutamide in pooled human liver microsomes (HLM) with or without 20 μM FABP1 WT or A94 variant. The formation rate of 4’-hydroxydiclofenac (4’OH-DCF) as a function of total **(A)** and unbound **(B)** diclofenac concentration. The formation rate of 4’-hydroxytolbutamide (4’OH-TBU) as a function of total **(C)** and unbound **(D)** tolbutamide concentration. In all panels, a representative experiment of three independent replicate experiments is shown. The points are individual measured velocities within the experiment, and solid lines are nonlinear least-squares fits of the Michaelis–Menten equation to the data. The kinetic parameters are summarized in Table 2. Data from additional replicates performed on separate days can be found at **Supplementary Figure 6**.

**Table 2.** Michaelis–Menten kinetic parameters for CYP2C9-mediated metabolism of unbound diclofenac and tolbutamide in human liver microsomes in the absence or presence of 20 μM FABP1 WT or A94 variants. Parameters are reported from model fits to unbound substrate concentration versus velocity data. Values are shown as mean ± standard deviation of three independent experiments. Each parameter was compared between no FABP1 and FABP1 WT, no FABP1 and FABP1 A94, and FABP1 WT and FABP1 A94 using ratio paired t-tests for each pairwise comparison.

|  | **Diclofenac** |  |  | **Tolbutamide** |  |  |
| --- | --- | --- | --- | --- | --- | --- |
|  | No FABP1 | +FABP1<br>WT | +FABP1 A94 | No FABP1 | +FABP1<br>WT | +FABP1 A94 |
| $V_{\max}$<br>(pmol/min/mg) | 770 ± 160 | 740 ± 170<sup>n.s.</sup> | 760 ± 180<sup>n.s.,n.s.</sup> | 280 ± 50 | 380 ± 60<sup>**</sup> | 350 ± 50<sup>n.s.,n.s.</sup> |
| $K_{m,u}$<br>(μM) | 7.4 ± 2.2 | 1.9 ± 0.8<sup>*</sup> | 3.1 ± 1.7<sup>*,*</sup> | 620 ± 60 | 260 ± 10<sup>*</sup> | 200 ± 30<sup>**,n.s.</sup> |
| $CL_{int,u}$<br>(μL/min/mg) | 110 ± 30 | 420 ± 90<sup>*</sup> | 270 ± 80<sup>*,*</sup> | 0.5 ± 0.1 | 1.5 ± 0.2<sup>*</sup> | 1.8 ± 0.3<sup>**,*</sup> |

The raw $p$ values were corrected for multiple comparisons by the Holm–Bonferroni procedure applied separately within each kinetic parameter, treating the three pairwise comparisons of a given parameter as one family of tests. Significance is reported at a family-wise error rate of $\alpha = 0.05$ . The superscript in the “+ FABP1 WT” column indicates the significance for the no FABP1 vs. FABP1 WT comparison. The superscripts in the “+ FABP1 A94” column indicate the significance for no FABP1 vs. FABP1 A94 comparison (first) and FABP1 WT vs. FABP1 A94 comparison (second). n.s., not significant; \*, adjusted $p$ value $< 0.05$ ; \*\*, adjusted $p$ value $< 0.01$ .

When tolbutamide was used as the CYP2C9 substrate in pooled HLMs, metabolite formation kinetics were affected differently by the presence of FABP1 than when diclofenac was used as a substrate. This shows substrate specific effects of FABP1 on drug metabolism. The Michaelis-Menten model fitted to total tolbutamide concentration versus product formation velocity data was significantly different when either FABP1 WT or A94 was present compared to absence of FABP1 (**Fig. 4C and Supplementary Table 3**). When unbound concentrations of tolbutamide were predicted in the presence of FABP1 as described above for diclofenac, the kinetic parameters that best fit the data were significantly different when FABP1 WT or A94 was present compared to no FABP1 controls (**Fig. 4D and Table 2**). This was due to significantly lower tolbutamide K_m,u_ in the presence of the FABP1 proteoforms and significantly higher V_max_ in the presence of WT FABP1 and a trend towards higher V_max_ in the presence of A94 FABP1. The calculated unbound intrinsic clearance increased significantly, 3.3- to 3.9-fold, in the presence of FABP1 (**Table 2**). The unbound intrinsic clearance for tolbutamide hydroxylation was also significantly different between FABP1 WT and A94.

The fact that the K_m,u_ and V_max_ were altered in the presence of FABP1 suggests that a more complex model than simple Michaelis-Menten kinetics may be needed to adequately describe the metabolism of diclofenac and tolbutamide in the presence of FABP1. Two models, adapted from a previously proposed substrate channeling model structure,^38,39^ were considered: model 1 in which unliganded FABP1 (apo-FABP1) does not interact with CYP2C9 (**Fig. 5A**) and model 2 in which apo-FABP1 interacts with CYP2C9 (**Fig. 5D**). For each model, unbound substrate concentrations, unbound FABP1 concentrations, and metabolite formation rates from pooled HLM experiments were used for parameter estimation. Both models incorporate channeling of the FABP1-bound substrate to CYP for metabolism. V_max_ and K_m,u_ for 4’OH-DCF and 4’OH-TBU formation were constrained to the estimated values from Michaelis-Menten fits from experiments done without FABP1 in parallel. For model 1, the parameters associated with substrate channeling (α, β) between FABP1 and CYP2C9 were estimated for 4’OH-DCF and 4’OH-TBU formation separately by simultaneous nonlinear least-squares regression of the model for each FABP1 proteoform (**Supplementary Fig. 7A-D)**. The possibility that apo-FABP1 also inhibits CYP2C9 via protein-protein interactions was assessed by fitting model 2 to the data as described above but assuming an additional kinetic parameter (K_i_) for apo-FABP1-CYP2C9 interaction (**Supplementary Fig. 7E-H**). For both drugs and both models, none of the fitted parameters were statistically different between the WT and A94 FABP1 based on extra sum-of-squares F-tests (**Supplementary Fig. 7I**).

**Figure 5.**
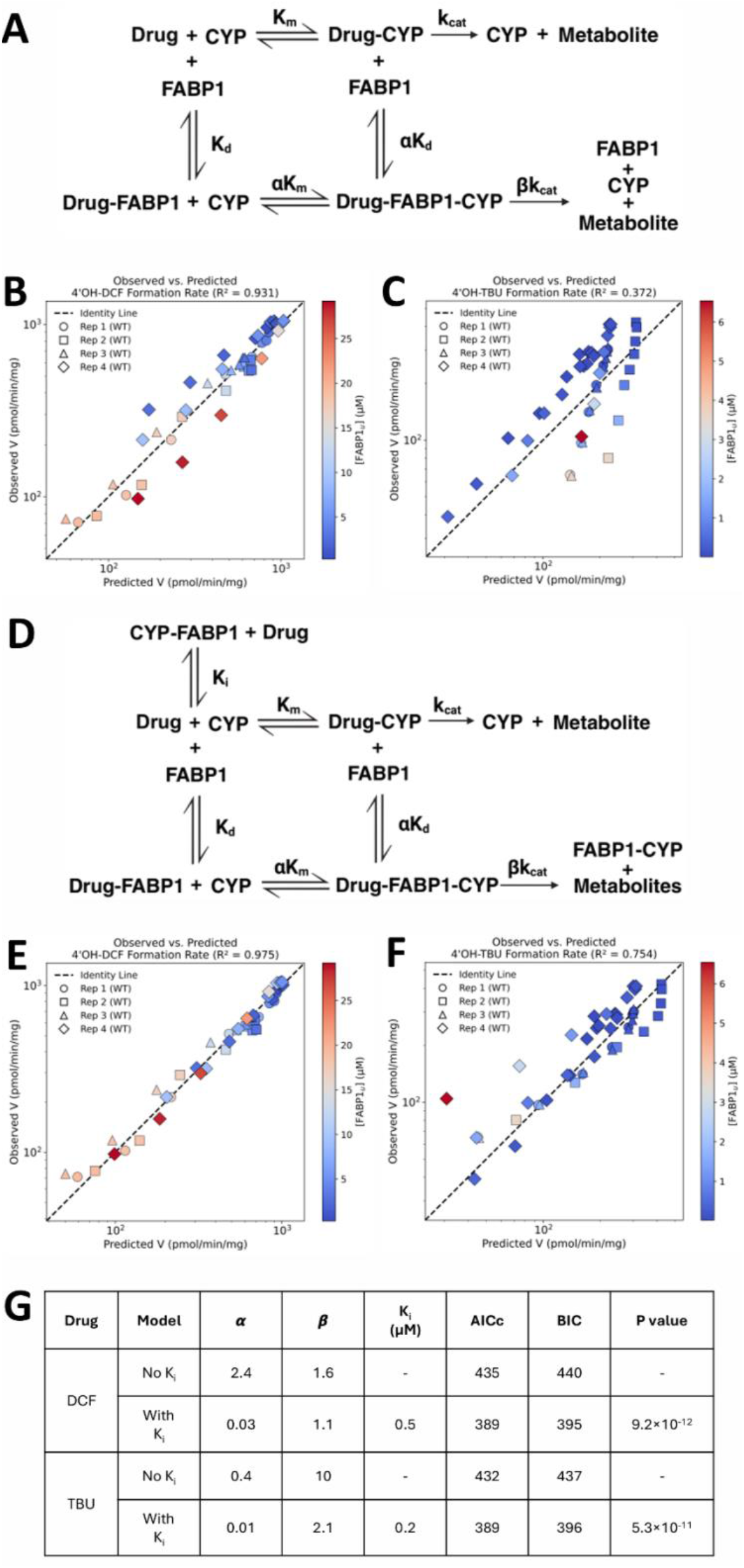
Estimation of kinetic parameters for diclofenac and tolbutamide metabolism with the different FABP1-CYP interaction models. **A** Kinetic scheme assuming apo-FABP1 does not interact with CYP2C9. **B-C** Observed versus predicted 4’OH-diclofenac (4’OH-DCF) or 4’OH-tolbutamide (4’OH-TBU) formation rate assuming the model in panel **A**. **D** Kinetic scheme assuming apo-FABP1 interacts with CYP2C9. **E-F** Observed versus predicted 4’OH-DCF and 4’OH-TBU formation rate assuming the model in panel **D**. **G** Summary table of best fitted α, β, and K_i_ assuming models in panels **A** and **D** for diclofenac and tolbutamide respectively. No K_i_ and with K_i_ models were compared using small-sample-corrected Akaike Information Criterion (AICc), Bayesian Information Criterion (BIC), and extra sum-of-squares F test for diclofenac and tolbutamide respectively. Different shapes represent the independent replicates completed on separate days with different batches of FABP1 purified. Color indicates the unbound FABP1 concentration. R^2^, coefficient of determination.

To better estimate the kinetic parameters for downstream *in vivo* clearance extrapolation, another replicate experiment was performed with varying concentrations of WT FABP1 (**Supplementary Fig. 8**). Formation of 4’OH-DCF and 4’OH-TBU was measured in pooled HLMs with three different total FABP1 concentrations resulting in distinct unbound substrate concentrations and a wider range of unbound FABP1 concentrations from the experiments in **Fig. 4** focused on understanding the WT and A94 differences. All kinetic data in **Fig. 4** and **Supplementary Fig. 8** were pooled for WT FABP1 and parameters were estimated with global fit of either model 1 or 2 (**Fig. 5**). Whether model 2 that includes apo-FABP1-CYP2C9 interaction was a better model to describe the data than model 1 was evaluated using small-sample-corrected Akaike information criterion (AICc), the Bayesian information criterion (BIC), and nested extra-sum-of-squares F-test. For both 4’OH-DCF and 4’OH-TBU formation data, model 2, the substrate channeling model with K_i_ term, yielded lower AICc and BIC, signifying better model fits. Furthermore, addition of K_i_ explained significantly more variance in 4’OH-DCF and 4’OH-TBU formation data than excluding the K_i_ (**Fig. 5G**). The resulting model fits yielded similar apo-FABP1-CYP2C9 K_i_-values (0.5 µM and 0.2 µM) with 4’OH-DCF and 4’OH-TBU formation data (**Fig. 5G)**.

### Incorporation of FABP1 improves prediction of metabolite formation in HLS9

Based on the observation that recombinant FABP1 alters the metabolite formation kinetics in HLMs, we hypothesized that native FABP1 in human liver cytosol will also impact metabolite formation by CYP2C9. To test this hypothesis, we incubated diclofenac and tolbutamide with the same set of HLS9 prepared from the UW human liver bank as described above (**Fig. 6A**). Data-independent acquisition mass spectrometry (DIA-MS) data was collected from the matched HLS9 samples. These data allow the simultaneous quantification of CYP2C9 plus multiple other proteins which may affect drug metabolism. Overall ∼89,000 precursors and ∼8,000 proteins were detected in these samples. The complete dataset was deposited on Panorama Public and is freely available for independent data analysis and further use (ProteomeXchange ID: PXD085173).

**Figure 6.**
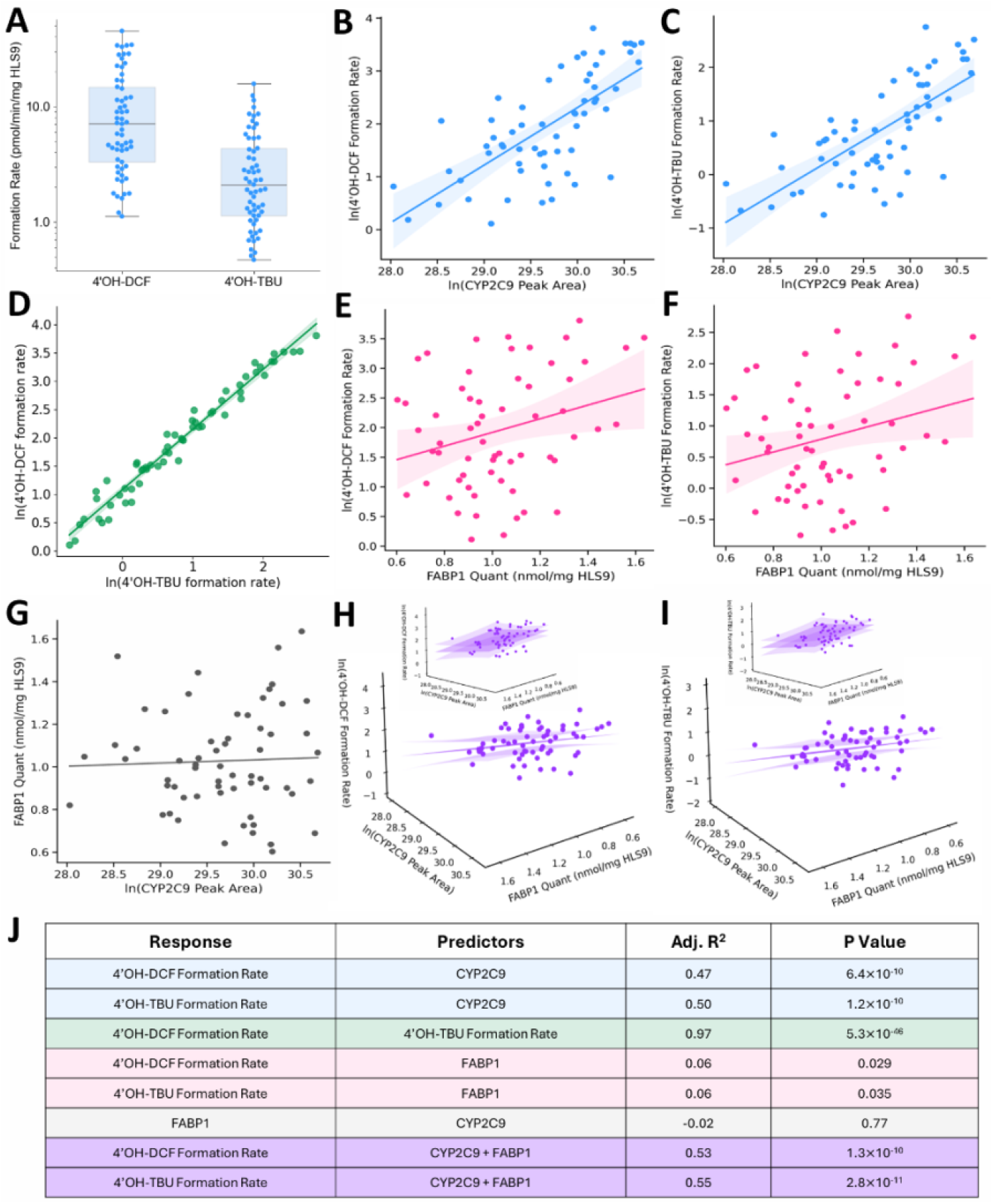
Relationships between CYP2C9 relative abundance, FABP1 expression level, and CYP2C9-mediated metabolite formation in human liver S9 fractions (HLS9). Solid lines are ordinary least-squares regression fits and shaded bands are the corresponding 95% confidence intervals in panels **B-I**. **A** Box and whisker plots of 4’OH-DCF and 4’OH-TBU formation rates in the 61 HLS9, plotted on log scale. **B-C** ln(4’OH-DCF formation rate) and ln(4’OH-TBU formation rate) as a function of CYP2C9 relative abundance. **D** Scatter plot of ln(4’OH-DCF formation rate) versus ln(4’OH-TBU formation rate). **E-F** ln(4’OH-DCF formation rate) and ln(4’OH-TBU formation rate) as a function of total FABP1 expression level. **G** Scatter plot of FABP1 expression versus ln(CYP2C9 peak area). **H-I** Three-dimensional regression surfaces relating metabolite formation to both predictors jointly, ln(CYP2C9 peak area) and FABP1 expression level for ln(4’OH-DCF formation rate) and ln(4’OH-TBU formation rate). The tangent line of the fitted regression surfaces are shown with 95% confidence interval surfaces on both sides. The insets show the regression surfaces at a different angle. **J** Summary table of the adjusted coefficient of determination (adj. R²) and p values for each regression fit in panels **B-I**. 4’OH-DCF, 4’hydroxydiclofenac; 4’OH-TBU, 4’hydroxytolbutamide.

The formation rates of 4’OH-DCF and 4’OH-TBU in the HLS9 correlated with relative CYP2C9 abundance measured from the DIA-MS data (**Fig. 6B,C**). The metabolite formation rates also correlated highly with each other (**Fig. 6D**) as expected for reactions carried out by the same metabolic enzyme. These results are consistent with CYP2C9 being the main enzyme hydroxylating diclofenac and tolbutamide. To further evaluate the effect of FABP1 on CYP2C9-mediated metabolism in HLS9, correlations between FABP1 expression quantified in the targeted assay (**Fig. 2**) and 4’OH-DCF or 4’OH-TBU formation rates were examined. The formation rates of 4’OH-DCF and 4’OH-TBU showed a weak positive correlation with FABP1 expression (**Fig. 6E,F**). Since FABP1 and CYP2C9 did not co-correlate (**Fig. 6G**), the positive correlation between FABP1 expression and tolbutamide and diclofenac hydroxylation suggests that 4’OH-DCF and 4’OH-TBU formation were higher with greater FABP1 concentration independent of CYP2C9 expression. This agrees with the HLM data where higher unbound intrinsic clearances were observed in the presence of FABP1 than in its absence.

Multiple linear regression models were built between 4’OH-DCF and 4’OH-TBU formation rates and relative abundance of CYP2C9 and total FABP1 or WT and A94 proteoforms of FABP1 as separate variables. Significant correlations between metabolite formation rates and FABP1 and CYP2C9 expression were observed (**Fig. 6H,I).** Model comparison metrics R^2^, adjusted R^2^, AICc, and BIC suggest that CYP2C9 relative expression level together with total FABP1 expression level predict the 4’OH-DCF and 4’OH-TBU formation rates the best (**Supplementary Table 4**). Statistically, F tests after Holm step-down multiple hypothesis correction for comparing these sets of nested models implies that the data provide enough evidence that the added FABP1 expression level, either as total FABP1 or separated by proteoforms, improves the model fit beyond random chance for 4’OH-DCF and 4’OH-TBU formation rates (**Supplementary Table 4**). However, BIC did not improve when WT and A94 FABP1 were assigned separate parameters (ΔBIC = 0), suggesting that the gain in fit did not justify modelling the two proteoforms separately (**Supplementary Table 4**). These findings suggest that native FABP1 in the human liver cytosol also alters CYP2C9 activity and support the model of FABP1-CYP2C9 interaction.

The acquired DIA-MS data allowed the relative quantification of several other proteins previously reported to affect CYP activity^12,40–42^: relative abundances of albumin (ALBU), cytochrome b5 (CYB5), NADH cytochrome b5 reductase (Cytb5R), glutathione S-transferase A1 (GSTA1), and cytochrome P450 oxidoreductase (POR) were quantified. The relative expression of CYP2C9 was found to positively correlate with the three ER-resident redox partners (CYB5, Cytb5R, POR) and with GSTA1, but not with albumin or FABP1 (**Supplementary Fig. 9**). Multiple linear regression models including CYP2C9, ALBU, CYB5, Cytb5R, GSTA1, POR, and FABP1 relative abundance post standardization to z-scores were built to test whether these other relevant proteins further explain variability in metabolite formation. In the presence of all the other proteins, only CYP2C9 and FABP1 had significant non-zero slopes (**Table 3**). Even when individual FABP1 WT and A94 expression levels were included, CYP2C9, FABP1 WT, and FABP1 A94 had significant contributions to predictions of the 4’OH-DCF and 4’OH-TBU formation rates (**Table 3**).

**Table 3.** Summary of coefficients and p values using the multiple linear regression predicting CYP2C9-mediated metabolite formation. Two outcomes were modeled, ln(4’OH-DCF formation rate) and ln(4’OH-TBU formation rate), with coefficients and p values reported side by side for each. Both models included a set of protein covariates as ln(peak area): CYP2C9, albumin (ALBU), cytochrome b5 (CYB5), NADH cytochrome b5 reductase (Cytb5R), glutathione S-transferase A1 (GSTA1), cytochrome P450 oxidoreductase (POR), and FABP1. All variables, including the response, were standardized to z-scores using the mean and standard deviation of the donors included in each model. Model 1 included total FABP1 expression as a single covariate, whereas Model 2 included FABP1 WT and A94 variant expression as separate covariates. 4’OH-DCF, 4-hydroxydiclofenac; 4’OH-TBU, 4’hydroxytolbutamide.

| Model | Covariates | 4'OH-DCF formation rate |  | 4'OH-TBU formation rate |  |
| --- | --- | --- | --- | --- | --- |
|  |  | Coefficient | P value | Coefficient | P value |
| 1 | CYP2C9 peak area | 0.58<br>[0.33, 0.84] | 0.00003 | 0.54<br>[0.30, 0.79] | 0.00004 |
|  | ALBU peak area | 0.08<br>[-0.12, 0.28] | 0.419 | 0.13<br>[-0.05, 0.32] | 0.156 |
|  | CYB5 peak area | 0.17<br>[-0.12, 0.45] | 0.252 | 0.25<br>[-0.02, 0.52] | 0.0671 |
|  | Cytb5R peak area | 0.07<br>[-0.17, 0.30] | 0.574 | 0.03<br>[-0.02, 0.52] | 0.775 |
|  | GSTA1 peak area | -0.07<br>[-0.29, 0.14] | 0.500 | -0.07<br>[-0.27, 0.14] | 0.526 |
|  | POR peak area | -0.01<br>[-0.27, 0.24] | 0.925 | -0.02<br>[-0.22, 0.26] | 0.881 |
|  | FABP1 expression | 0.29<br>[0.10, 0.48] | 0.0033 | 0.29<br>[0.10, 0.48] | 0.0020 |
| 2 | CYP2C9 peak area | 0.59<br>[0.33, 0.85] | 0.00003 | 0.55<br>[0.30, 0.79] | 0.000040 |
|  | ALBU peak area | 0.08<br>[-0.12, 0.28] | 0.436 | 0.13<br>[-0.06, 0.32] | 0.165 |
|  | CYB5 peak area | 0.18<br>[-0.11, 0.47] | 0.228 | 0.26<br>[-0.01, 0.54] | 0.0599 |
|  | Cytb5R peak area | 0.06<br>[-0.17, 0.30] | 0.593 | 0.03<br>[-0.19, 0.25] | 0.795 |
|  | GSTA1 peak area | -0.08<br>[-0.30, 0.14] | 0.477 | -0.07<br>[-0.28, 0.14] | 0.503 |
|  | POR peak area | -0.01<br>[-0.27, 0.25] | 0.936 | 0.02<br>[-0.22, 0.26] | 0.872 |
|  | FABP1 WT expression | 0.43<br>[0.15, 0.70] | 0.0028 | 0.42<br>[0.17, 0.68] | 0.0017 |
|  | FABP1 A94 expression | 0.34<br>[0.07, 0.61] | 0.014 | 0.34<br>[0.09, 0.60] | 0.0090 |

### Simulation of hepatic clearance in the presence of FABP1 using minimal PBPK models

To predict the impact of endogenous FABP1 on CYP2C9-mediated clearance based on the *in vitro* data generated in this study, three minimal PBPK models with the same liver physiology and an average intracellular FABP1 expression level of 399 µM^10^ were developed: 1) with only drug-FABP1 binding affecting the liver distribution of the drugs, 2) with FABP1 binding and drug-FABP1-CYP2C9 interaction but without apo-FABP1-CYP2C9 interaction, and 3) with FABP1 binding and drug-FABP1-CYP2C9 interaction as well as apo-FABP1-CYP2C9 interaction (**Figure 7A**). According to the literature, *in vivo* 4’OH-DCF and 4’OH-TBU formation clearances are between 7.9 and 10.5 L/hr and 0.62 and 0.72 L/hr in healthy human population, respectively.^43–53^ Using the model with only drug-FABP1 binding and FABP1 only affecting drug distribution into the liver, the *in vivo* 4’OH-DCF and 4’OH-TBU formation clearances were predicted to be 1.5 L/hr and 0.07 L/hr (**Fig. 7B-C black bars**). The observed values were underpredicted by 81% and 89%. When channeling of drug from FABP1 to CYP2C9 was considered but without apo-FABP1-CYP2C9 interaction, the *in vivo* 4’OH-DCF and 4’OH-TBU formation clearances were predicted to be 27 L/hr and 33 L/hr respectively (**Fig. 7B-C pink bars**). This second model resulted in 157% and 4483% overprediction of the metabolite formation clearances. When the inhibitory interaction between apo-FABP1 and CYP2C9 was incorporated into the liver model, the *in vivo* 4’OH-DCF and 4’OH-TBU formation clearances were predicted to be 4.7 L/hr and 0.37 L/hr (**Fig. 7B-C, green bars)**, which were both within 2-fold of the observed values. The third model improved the clearance prediction by 3.1-fold and 5.3-fold for 4’OH-DCF and 4’OH-TBU formation *in vivo* when compared to the model without any FABP1-CYP2C9 interaction. As such the prediction incorporating both substrate channeling and apo-FABP1-CYP interactions most accurately predicted the *in vivo* clearance among the three liver models developed (**Fig. 7D**).

**Figure 7.**
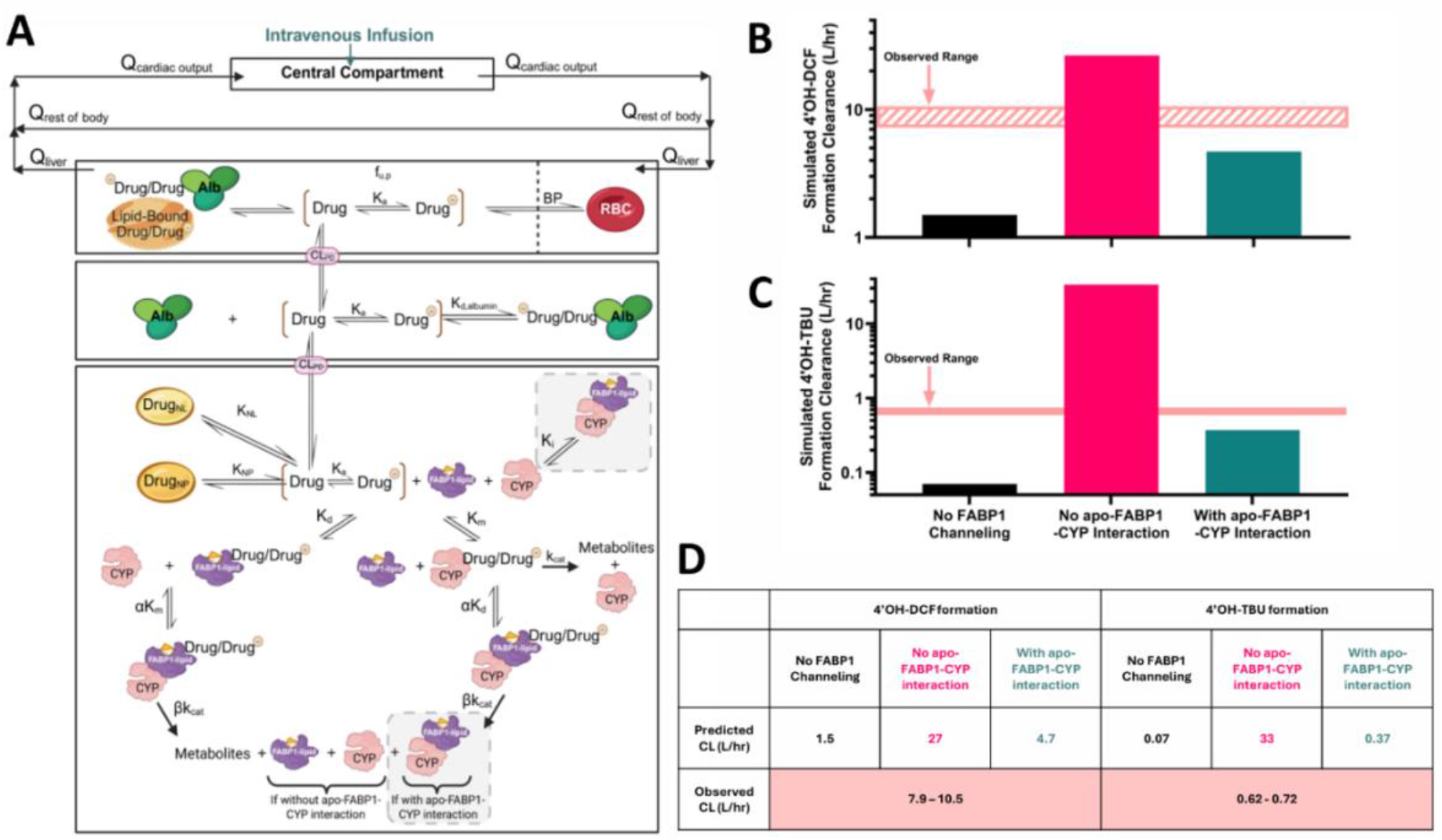
Prediction of hepatic formation clearance of 4’OH-diclofenac and 4’OH-tolbutamide using a minimal physiologically based pharmacokinetics (PBPK) model. A Structure of the physiologically based liver model incorporating different clearance schemes for clearance prediction. Q_cardiac output_, blood flow to the central compartment; Q_rest of the body_, blood flow to the rest of the body; Q_liver_, blood flow to the liver; CL_PD_, passive diffusion clearance; K_a_, acid dissociation constant; f_u,p_, unbound fraction of drug in the plasma; K_d,albumin_, dissociation constant between albumin and drug; K_NL_, partition constant for neutral lipid binding; Drug_NL_, drug partitioned into the neutral lipid; K_NP_, partition constant for neutral phospholipid binding; Drug_NP_, drug partitioned into the neutral phospholipid; K_d,FABP1_, dissociation constant between FABP1 and drug; K_i_, dissociation constant for the CYP-FABP1 complex; k_cat_, the turnover number; K_m_, the Michaelis constant; α and β are constants from the no apo-FABP1-CYP interaction model 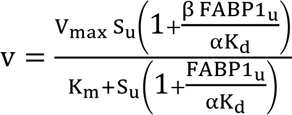 (without the grey boxed scheme) or with apo-FABP1-CYP interaction model 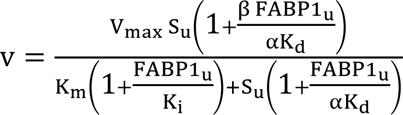 in **Figure 5**. B Comparisons of the predicted hepatic formation clearances of 4’OH-DCF assuming FABP1 has no impact on metabolism (No FABP1 Channeling), no apo-FABP1-CYP interaction model, and with apo-FABP1-CYP interaction model to observed *in vivo* clearances. C Comparisons of the predicted hepatic formation clearances of 4’OH-TBU assuming FABP1 has no impact on metabolism (No FABP1 Channeling), no apo-FABP1-CYP interaction model, and with apo-FABP1-CYP interaction model to observed *in vivo* clearances. The shaded area represents the literature reported range of formation clearance of 4’OH-DCF or 4’OH-TBU in healthy adults.^43–53^ D Summary of the predicted and observed clearance values in panel B-C. Panel A was created in BioRender. Wen, W. (2026) https://BioRender.com/w7204g9.

## Discussion

Accurate prediction of drug clearance is crucial in drug discovery and development.^54,55^ Poor predictions can result in failure or discontinuation of development of valuable compounds. Further, incorrect identification of the specific enzymes clearing the drug can result in inaccurate risk assessment of drug interactions putting patients at risk. *In vitro* incubations with human liver subcellular fractions are the cornerstone of clearance predictions.^56^ Yet, predictions often lack sufficient accuracy and fail to meet the general two-fold criterion for successful prediction. The poor extrapolation was also illustrated here with diclofenac and tolbutamide. The predicted *in vivo* clearances in the absence of FABP1 were 81% and 89% lower than the observed *in vivo* clearance. Collectively, our findings using diclofenac and tolbutamide as model drugs suggest that this failure in clearance predictions is at least in part due to ignoring cellular complexity and protein-protein interactions between FABP1 and drugs and CYPs.

The current study provides the first published evidence that quantitative prediction of CYP2C9-mediated metabolic clearance is improved by including FABP1 in the *in vitro* experiments and by incorporating native liver FABP1 expression into the *in silico* prediction methods. Since cytosolic contents are absent in HLMs, the impact of FABP1 on kinetics of CYP2C9 mediated 4’OH-DCF and 4’OH-TBU formation was examined in pooled HLMs with addition of recombinant FABP1. The changes in metabolite formation kinetics after addition of FABP1 cannot be explained by substrate sequestration to FABP1 or the free-drug hypothesis alone (**Fig. 4** and **Table 2**). Instead, the impact of FABP1 on metabolite formation kinetics is consistent with protein-protein interaction between FABP1 and CYPs similar to prior reports with recombinant CYP2C9 and CYP3A4.^12,16^ The increase in V_max_ of 4’OH-TBU formation in the presence of FABP1 suggests FABP1 increases CYP2C9 activity *in vitro* via substrate channeling from FABP1 to CYP2C9 (**Fig. 4** and **Table 2**). The increase in V_max_ is unlikely to be due to FABP1 removing inhibitory fatty acids from CYP2C9 active site as that mechanism would be expected to impact each substrate similarly and be a substate independent effect. However, the difference in the metabolite formation kinetics and kinetic parameters for 4’OH-DCF and 4’OH-TBU formation in the presence of FABP1 show that the effect of FABP1 is substrate dependent (**Fig. 5**). The interpretation of protein-protein interactions is in agreement with the literature reports of intracellular lipid binding protein-CYP interactions that include both substrate channeling and apo-CRABP or apo-FABP1 acting as a noncompetitive inhibitors of the CYP.^12,38^ The kinetic data obtained here suggest that ligand binding to FABP1 is not necessary for the FABP1-CYP interaction. The best fit model was the one incorporating inhibitory apo-FABP1-CYP2C9 interactions as well. The estimated K_i_ values for the apo-FABP1-CYP2C9 interaction in our study were in the nanomolar range. These K_i_ values are similar in magnitude to the previously reported apo-FABP1-CYP3A4 interaction that showed 15.3 nM K_i_.^12^

Human liver S9 fractions include cytosol and the native liver FABP1. Hence, we tested whether native FABP1 expression in HLS9 correlates with metabolite formation *in vitro*. Statistical modeling between CYP2C9 metabolic activity and expression of proteins contributing to drug metabolism measured via LC-MS/MS based proteomics suggests that CYP2C9 and FABP1 expression together explain majority of the variance observed between donors in metabolite formation rates. The relative expression of ALBU, CYB5, Cytb5R, GSTA1, and POR, which have previously been implicated as contributors to variability in drug metabolism kinetics,^12,40–42^ did not explain variability in diclofenac or tolbutamide metabolism in this study (**Table 3**). The estimated coefficient associated with FABP1 was positive, consistent with the facilitation of drug metabolism by FABP1 shown with the recombinant FABP1 (**Table 3**).

To define the quantitative impact of FABP1-CYP2C9 interactions in *in vitro* to *in vivo* clearance predictions we expanded our liver PBPK model^10^ to incorporate the FABP1-CYP2C9 substrate channeling schemes. The kinetic parameters estimated in pooled HLM data in the presence of recombinant FABP1 were incorporated into the model together with the known FABP1 expression in human liver. The values for FABP1 expression included in the model better reflect the human liver concentrations than the concentrations present in the experiments in HLS9 as HLS9 are diluted during homogenization and further in drug incubations resulting in diluted FABP1 concentration. The minimal PBPK model that incorporated apo-FABP1-CYP2C9 interaction together with the substrate channeling offered the most accurate, within 2-fold, prediction of the *in vivo* metabolite formation clearance. The predictions conducted without consideration of FABP1 in the liver underpredicted the *in vivo* tolbutamide and diclofenac clearance by 81-89% while the simulations conducted without incorporating apo-FABP1-CYP2C9 interaction overpredicted the *in vivo* clearance by 157-4483% (**Fig. 7**). The PBPK modeling results suggest apo-FABP1-CYP2C9 interaction is relevant *in vivo* and demonstrate the importance of taking FABP1 into account for more accurate IVIVE. Further the PBPK model developed provides a framework for future *in vitro* to *in vivo* clearance predictions.

The T to C SNP in exon 3 of the human FABP1 gene is highly prevalent.^17–20^ The resulting amino acid change has been associated with altered nuclear channeling of FABP1 and altered interactions of FABP1 with nuclear receptors such as PPARα and hepatocyte nuclear factor-4α (HNF4α), and with altered fatty acid processing in the liver.^17,57–61^ In our liver bank, the FABP1 expression was not different between FABP1 T94A genotypes, and FABP1 WT and A94 variants appeared equally expressed in T/C heterozygous individual donor livers (**Fig. 2**). In contrast, total FABP1 expression was significantly lower in fatty human livers (**Fig. 2**). This is in agreement with previous data of decreased FABP1 mRNA expression in human livers with MAFLD.^62^ In the UW human liver bank, carriers of C allele (T/C or C/C) appear to have elevated risk of MAFLD (odds ratio = 2.08) but the sample set is underpowered to reach statistical significance. This trend, however, aligns with the finding from 1106 Chinese study participants that reported a significant association between MAFLD risk and the C allele.^36^

The observation of clinical dyslipidemia and elevated risk of MAFLD associated with FABP1 T94A variant led us to hypothesize that FABP1 polymorphisms may alter ligand distribution into the liver.^36,62^ Previously, we reported that incorporation of FABP1 into PBPK model improves the prediction of liver K_p_ in humans.^10^ However, PBPK modeling with FABP1 WT and A94 variants incorporated separately here showed a low risk of altered tissue K_p_ in A94 expressors for the drugs modeled, likely due to the similar binding affinities of the drugs of interest between the two proteoforms (**Fig. 3 A-D**). Even though many studies have reported minimal effects of the T94A substitution on binding affinity for a broad variety of ligands,^17,19,20,57^ some endogenous ligands like endocannabinoids have been reported to have weaker binding affinity to the FABP1 A94 variant.^63^ Our model suggests that tissue K_p_ and cellular distribution of FABP1 ligands can be substantially lower in A94 expressors if binding affinity of the ligand to FABP1 is significantly weaker. A lower tissue distribution of lipids or drugs in A94 carriers is important as it is also predicted to result in lower cytosolic drug-FABP1 complex concentrations. Many drugs, like fenofibrates, activate PPARα to regulate lipid metabolism, triglycerides, and cholesterol levels.^64^ These drugs also bind FABP1 and are likely delivered to their target in the nucleus by FABP1. Hence, the variability in drug-FABP1 bound concentrations offers potential explanation for the observed reduced effects of fenofibrates in FABP1 A94 expressors in humans.^65^ The PBPK models presented here can also be adapted to incorporate FABP1-nuclear receptor interactions to provide insights on pharmacodynamic effects and FABP1 associated disease progression.

In conclusion, the metabolite formation data in pooled HLMs with recombinant FABP1 and 61 HLS9 collectively show that FABP1 facilitates CYP2C9-mediated metabolism in a substrate specific manner. We demonstrate that incorporating FABP1 into a PBPK model and to *in vitro-*to-*in vivo* clearance predictions improves the prediction accuracy for diclofenac and tolbutamide allowing predictions within 2-fold of observed values. These findings suggest that incorporation of FABP1 into clearance prediction workflows is important for accurate predictions. In general, researchers should screen for drug-FABP1 binding and consider integrating FABP1-CYP interaction into PBPK models to improve prediction accuracy. As FABP1 is also expressed in the kidney and intestines in humans, FABP1-CYP interactions can affect clearance prediction beyond liver. The dynamic models developed here can be adapted to other FABP1 expressing organs or be incorporated into whole body PBPK models and used to predict drug disposition. The minimal PBPK models developed can also be used and expanded to simulate lipid disposition for FABP1 WT and A94 expressors.

## Methods

### Expression, purification, and characterization of human FABP1 WT and A94 variants

The details of chemicals and reagents used in this study can be found in the **Supplementary Method I**. The Hexa-histidine-tagged A94 variant human FABP1 was cloned into a pET28a1 expression vector (**Supplementary Method II, Supplementary Fig. 11**) and expressed in Rosetta 2 *Escherichia coli* as described in **Supplementary Method II**. FABP1 was purified and delipidated as previously described^10,16^ (**Supplementary Method III, Supplementary Fig. 12)**. In brief, the his-tagged WT and A94 variant recombinant FABP1 were purified using nickel-nitrilotriacetic acid (Ni-NTA) affinity chromatography followed by size-exclusion chromatography yielding pure protein (∼ 40mg/L culture) with molecular weight of 14.5 kDa. The protein concentration was quantified via BCA (Pierce, Waltham, MA) assay and was stored in 50 mM potassium phosphate (KPi) buffer pH 7.4, with 100 mM KCl as previously described.^10^ The WT and A94 proteins were characterized using fluorescence spectroscopy and Far-UV circular dichroism (CD) (**Supplementary Method IV)** to evaluate ligand binding and conformational stability.

### Characterization of drug binding to FABP1 WT and A94 using fluorescence spectroscopy

The binding of drug ligands to WT and A94 FABP1 was estimated using fluorescent ligand displacement assay as previously described.^66^ Briefly, the fluorescent probe DAUDA (0.5 µM) was first pre-bound with FABP1 (0.3 µM) WT or A94 in assay buffer (50 mM KPi, 100 mM KCl, pH 7.4) in a clear quartz cuvette. Methanolic solutions of diclofenac (0.2–127 µM), gemfibrozil (0.2–84 µM), pioglitazone (0.2–7.9 µM), and tolbutamide (0.3–132 µM) were added in 12 points. All ligands tested were confirmed to lack background fluorescence at 30 µM concentration in buffer or in the presence of 0.3 µM FABP1. A ternary complex model was assumed when attenuation of blue-shifted fluorescence traces was observed after saturating concentrations of ligands were added.^16^ Specific fluorescence was acquired from singular value decomposition (SVD) analysis and fitting was completed in COPASI as previously described.^16^ K_d_ values are reported (± standard deviation) from three independent replicate data sets generated on different days with different batches of FABP1 WT or A94 proteins purified and done paired on each day.

### Human liver bank and preparation of S9 fractions

A total of 61 human liver samples from the University of Washington Human liver Bank^34,35^ were included in the study. Demographic information for liver tissue donors and liver pathology can be found in **Supplementary Table 2**. Human liver S9 fractions (HLS9) were prepared as previously published ^34^. Briefly, 151.8–202.5 mg of tissue (except two donors with 46.8 mg and 51.7 mg of tissue) was cut and transferred into the tissue homogenizing tubes with ceramic beads. A constant ratio (5-fold v/w) of homogenizing buffer (50 mM KPi, 250 mM sucrose, 50 mM KCl, pH 7.4 with EDTA-free protease inhibitor cocktail tablet) to liver tissue was added directly to the tissue homogenizing tubes prior to homogenization. The tissue was then homogenized using Omni Bead Ruptor 24 (Kennesaw, GA) using five 20-second disruption cycles at 5.00 m/s with a 3-second rest between cycles between -15 °C to -10 °C, followed by centrifugation at 9,000 g and storage in homogenizing buffer as previously reported.^10^

### DNA extraction from human liver samples and FABP1 promoter and T94A genotyping

Genomic DNA was extracted from human liver tissues using the QIAamp DNA Mini kit (Qiagen, Germantown MD, Cat# 51304) according to manufacturer’s instructions for tissues and as previously described.^67^ DNA concentration was measured by NanoDrop at 260 nm (Thermo Fisher Scientific, Waltham, MA), and purity estimated from A260/280 ratio. DNA samples were stored at 4°C for genotyping assays. FABP1 promoter (Thermo Fisher, Waltham, MA; rs2919872 for C>T) and T94A (Thermo Fisher, Waltham, MA; rs2241883 for T>C) alleles were genotyped using StepOnePlus instrument (Thermo Fisher Scientific, Waltham, MA). Reactions were run using the manufacturer’s standard genotyping protocol. Genotypes were assigned using StepOnePlus software from the allelic discrimination plot.

### Quantification of FABP1 WT and A94 in HLS9 Fractions Using UHPLC-MS/MS

FABP1 expression in the 61 individual donors was previously measured.^10^ To estimate the fraction of total FABP1 that is present as WT and A94 proteoforms, the 61 HLS9 were analyzed using liquid chromatography-tandem mass spectrometry (LC-MS/MS) and measuring a surrogate peptide for the polymorphic residue (LVTTFK for WT and LVTAFK for A94 FABP1; **Supplementary Table 4**). Analysis was done using a Sciex 5500 QTrap mass spectrometer (Sciex) coupled to an Agilent 1290 LC (Agilent). Peptides were separated using an Aeris peptide XB-C18 column (50 ×2.1 mm, 1.7 μm), and a SecurityGuard Ultra LC C18-peptide cartridge (Phenomenex). Tissue and protein digestion and sample preparation, gradient elution, and peak integration were done as previously described.^10^ For quantification, the ratio of LVTTFK or LVTAFK peptide peak area to peptide area of 20 nM F[^13^C_9_ ^15^N]TITAGSK was used to construct standard curves and calculate concentrations of each FABP1 proteoform in individual HLS9. Briefly, calibration curves and quality control (QC) samples for FABP1 quantification were prepared by spiking purified human FABP1 (WT and A94 mixed in 1:1 ratio) into 0.2 mg/mL mouse liver S9 (final concentration 25—500 nM for each proteoform for the standard curve and 25, 60, 160, and 270 nM for lower limit of quantitation, low QC, medium QC, and high QC). The average of the three independent singlet experiments done on 3 separate days is reported.

### Simulation of liver tissue-to-plasma partition coefficient (K_p_) and intracellular drug-FABP1 concentration

A previously published^10^ physiologically based liver distribution model with human physiology and drug-FABP1 binding was used to predict drug distribution to the liver. The original model was modified to include FABP1 WT and A94 proteoforms separately, allowing for input of expression levels and drug-binding affinity for each proteoform quantified in this study. The updated model is available at GitHub: Isoherranen-Lab. The liver distribution of diclofenac, pioglitazone, gemfibrozil, and tolbutamide in the 61 human livers was predicted using the measured K_d_ values and individual FABP1 WT and A94 expression levels. The predicted liver K_p_ and cytosolic drug-FABP1 concentration were output as previously reported.^10^

### Kinetics of 4’-OH-DCF and 4’-OH-TBU formation in the presence and absence of FABP1 in pooled HLMs

The kinetics of 4’OH-DCF and 4’-OH-TBU formation in the presence and absence of FABP1 in pooled HLMs were determined under conditions of protein and time linearity. Experiments with and without FABP1 were done as matched pairs on the same day with technical duplicates. HLMs pooled from 50 donors were purchased from Gibco (Cat. HMMCPL, Thermo Fisher Scientific, Waltham, MA). Pooled HLMs (0.01 mg/mL protein) with or without 20 µM WT or A94 FABP1 were preincubated with eight different concentrations of diclofenac (0.9 – 50 µM) or tolbutamide (20 – 1600 µM) for 5 minutes at 37 °C in 45 µL incubation buffer (50 mM KPi, pH 7.4, 100 mM KCl) in a 96-well plate. Reactions were initiated by adding NADPH (1 mM final concentration in 50 µL incubation) and quenched after 10 minutes with 50 µL of ice-cold acetonitrile containing 25 nM 4’OH-5-chloro-diclofenac or 10 nM 4’OH-mephenytoin as internal standards for 4’OH-diclofenac and 4’-OH-tolbutamide assays respectively. The quenched reactions were centrifuged at 3,200 g for 60 minutes at 4 °C and 60 µL supernatant was transferred to 96-well MS plate for LC-MS/MS analysis. The concentrations of 4’OH-DCF and 4’OH-TBU were analyzed either using a Sciex API 5500 or Sciex API 6500+ QTRAP mass spectrometer (Sciex) coupled to an Agilent 1290 or Agilent 1290 Infinity II Ultra-High-Performance Liquid Chromatography system (Agilent) as previously published.^14^ The details of the bioanalytical methods are described in **Supplementary Methods V**.

Free concentrations of diclofenac and tolbutamide in the HLM incubations in the presence of FABP1 were calculated as previously reported^15^ using the drug-FABP1 binding affinity, unbound fraction of drug in HLM, and total FABP1 concentration in the incubations. The Michaelis-Menten model was fit to the 4’OH-DCF and 4’OH-TBU formation data in GraphPad Prism 10 using nominal and free concentrations of diclofenac and tolbutamide, to determine the apparent and unbound metabolite formation kinetics. K_m_ and V_max_ values are reported as arithmetic means ± standard deviation from experiments done on three separate days. For every replicate experiment, the model fits were compared in GraphPad Prism using the nonlinear regression comparison and F test as previously described.^16^ Paired t tests were used to evaluate whether K_m_, V_max_, or CL_int_ (calculated as the ratio of V_max_ and K_m_) were different between two groups. Holm-Bonferroni step-down method was used to account for multiple hypothesis testing. Significance was assessed at a predetermined α level of 0.05.

### Estimation and verification of kinetic parameters using metabolite formation data in the pooled HLMs

The previously described^38^ substrate channeling models in which either only drug bound FABP1 interacts with CYP2C9 without apo-FABP1-CYP2C9 interaction (equation 1) or in which both liganded and unliganded FABP1 interact with CYP2C9 (equation 2) were fit to the metabolite formation rate versus unbound substrate and FABP1 concentration data:

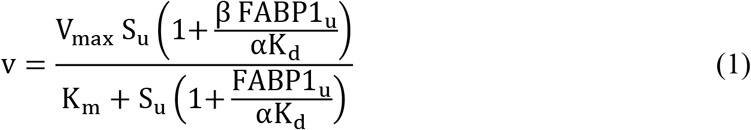

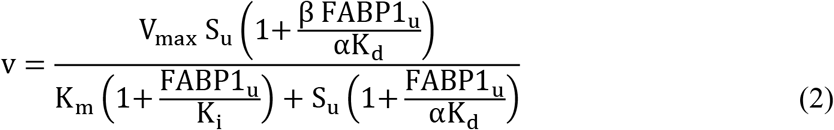

In these equations, S_u_ is the unbound substrate concentration and v is the formation rate of the product (4’OH-DCF or 4’OH-TBU). For each replicate, the constants K_m_, V_max_, and the drug-FABP1 binding affinity K_d_ were fixed to the values independently determined, while two (α and β) or three parameters (α, β, and K_i_) were estimated using equation 1 and 2, respectively. Parameters were estimated separately for WT and A94 FABP1 by complete pooling: a single set of α and β for equation 1 or a single set of α, β, and K_i_ was fit jointly to all replicates of a variant by minimizing the pooled residual sum of squares, with each replicate contributing its own fixed constants. Optimization used bounded nonlinear least squares under ordinary least squares, with parameters constrained to biologically plausible ranges ( α ∈ [0.01,10], β ∈ [0,10], K_i_ ∈ [0.01,10]). WT and A94 FABP1 effects were compared by an extra sum-of-squares F test comparing a model with all parameters shared between the two proteoforms against one with proteoform-specific parameters. Another independent experiment was performed to measure the velocity of 4’OH-DCF and 4’OH-TBU formation at different substrate and WT FABP1 concentrations as described in **Supplementary Method VI**. The WT FABP1 data were pooled together to estimate the kinetic parameters as described above for *in vivo* clearance extrapolation. All analyses were performed in Python (version 3.11.9) using NumPy, SciPy, and pandas. Raw data and code can be found at GitHub: Isoherranen-Lab.

### Formation rates of 4’OH-DCF and 4’OH-TBU in a panel of 61 HLS9

The formation of 4’OH-DCF and 4’OH-TBU was measured in the panel of 61 HLS9. All HLS9 experiments were performed in duplicates at 0.2 mg/mL S9 protein in 100 mM KPi, pH 7.4 in 96-well plates and repeated on two separate days. After a 6-minute pre-incubation at 37 °C in the presence of substrate (2 μM diclofenac or 50 μM tolbutamide), reactions were initiated with NADPH (1 mM final concentration) and allowed to proceed for 30 min for tolbutamide and 7 min for diclofenac at 37 °C in a water bath with a total reaction volume of 100 μL. The reactions were quenched with an equal volume of acetonitrile containing internal standard and prepared and analyzed via LC-MS/MS as described above.

### Global proteomics by data-independent acquisition mass spectrometry (DIA-MS) and statistical modeling between protein expression and metabolite formation data in HLS9

Complete details on protein digestion, mass spectrometry, and data processing are provided in **Supplementary Method VII**. The relationship between 4’OH-DCF and 4’OH-TBU formation rate and proteomic covariates was assessed by ordinary least squares (OLS) multiple linear regression and modeled on the natural-logarithm scale. For 4’OH-DCF and 4’OH-TBU formation rate, CYP2C9 and FABP1 abundances were evaluated using nested models: CYP2C9 alone (Model 1), addition of total FABP1 (Model 2), and both variants additively (Model 3). All models for a given outcome used the same observations to allow valid information-criterion comparisons.

To further evaluate the associations between metabolite formation rates and a list of drug metabolism related proteins that have been reported to affect CYP activities and clearance prediction,^12,40–42^ two multivariable models were fit to predict 4’OH-DCF and 4’OH-TBU formation rate respectively. Model 1 included the relative abundances of CYP2C9, ALBU, CYB5, Cytb5R, GSTA1, POR, and expression level of FABP1 as predictors. Model 2 was an alternative parameterization in which total FABP1 was replaced by the separately quantified WT and A94 FABP1, with the remaining predictors unchanged. For each coefficient, the null hypothesis that the coefficient equals zero was tested using two-sided t-test, and the resulting p-value was used to determine whether that covariate was independently associated with activity.

Models were compared using AICc and BIC. AICc was used because of the modest number of observations (n=61). The model comparison metrics were compared for 4’OH-DCF and 4’OH-TBU outcomes separately. Each model was also compared against the base model by a nested extra-sum-of-squares F-test. Within each outcome, the resulting p-values were corrected for multiple comparisons by the Holm step-down procedure, which controls the family-wise error rate. Significance was assessed at a predetermined α level of 0.05. All analyses were performed in Python (version 3.11.9) using statsmodels, with scipy, numpy, pandas, and matplotlib. The raw data and code are available at GitHub: Isoherranen-Lab.

### Prediction of metabolite formation clearance *in vivo* using minimal PBPK model incorporating FABP1 effects

Adapted from the previously established liver distribution model,^10^ three different liver models were developed in MATLAB & Simulink (R2025b) to predict the impact of FABP1 on CYP2C9-mediated clearance with the same liver physiology: 1) with only drug-FABP1 binding, 2) with drug-FABP1-CYP2C9 interaction but without apo-FABP1-CYP2C9 interaction, and 3) with drug-FABP1-CYP2C9 interaction as well as apo-FABP1-CYP2C9 interaction (**Figure 7A**). The developed liver models are published at GitHub: Isoherranen-Lab. Metabolism was assumed to occur exclusively intracellularly. The CYP-mediated Michaelis-Menten kinetics or substrate channeling enzyme kinetic scheme was incorporated directly into the intracellular compartment. Parameters (Model 1: K_m_, V_max_, and K_d_; Model 2: K_m_, V_max_, K_d_, α, and β; Model 3: K_m_, V_max_, K_d_, α, β, and K_i_) estimated by fluorescence titration and nonlinear regression of the *in vitro* metabolite formation rate data as described above were used in the simulation. Intracellular FABP1 expression level of 399 µM was used as input value for the simulations as the previously measured mean concentration of FABP1 in the liver.^10^ Using a reported CYP2C9 expression level of 127 pmol/mg microsomal protein,^78^ the k_cat_ for 4’OH-DCF and 4’OH-TBU formations in the whole liver were predicted to be 463 hr^-1^ and 116 hr^-1^ respectively using the *in vitro* V_max_ (in pmol metabolite/min/mg microsomal protein) data generated in this study. Intracellular CYP2C9 concentration was calculated to be 9 µM taking the CYP2C9 expression level,^78^ an average microsomal protein yield of 40 mg/g liver,^79^ and fraction of liver wet weight as intracellular water^10,80^ into account. The governing equations and physiological parameters for a 70 kg human^10,81^ are listed in **Supplementary Method VIII** and **Supplementary Table 6-7**.

The model was solved for a constant-rate intravenous infusion into the systemic compartment at infusion rate under linear condition, with all compartments initialized at zero concentration. Simulations were run for 200 hours, sufficient for all compartments to reach steady state. Attainment of steady state was verified visually and numerically. At steady state, clearance was calculated as the ratio of infusion rate and the steady-state plasma concentration C_ss_. *In vivo* formation clearance of 4’OH-DCF and 4’OH-TBU in healthy human population were retrieved from the literature. Simulated clearance using Models 1-3 with different kinetic schemes assumed were compared to the average of the observed *in vivo* clearance as a fold error (ratio of simulated and observed clearance). The model with the least fold error is determined to be the best liver model for IVIVE in this study.

## Supporting information

Supplemental results and materials

## Data and code availability

Raw data and code for the statistical analysis and reproducing the figures can be found at GitHub: Isoherranen-Lab. The targeted MRM quantification of FABP1 along with the entire human liver DIA dataset can be found on Panorama using the URL: https://panoramaweb.org/uw-human-liver-bank-s9.url) and were assigned the ProteomeXchange^77^ ID: PXD085173.

## Acknowledgements

We would like to thank Martin Sadilek and Brandon Bol for their training on the circular dichroism spectroscopy.

## Author contributions

Y.W.W. performed statistical and physiologically based pharmacokinetic modeling, protein expression and purification, protein characterization, fluorescence titrations, human liver S9 fraction preparation, targeted FABP1 quantification, and drug incubations. E.F. performed fluorescence titrations, DNA extraction and genotyping, and drug incubations, using protocols provided by Y.W.W., Y.W.W., N.I., and A.N. designed and Y.W.W. conducted and analyzed the stopped flow and CD spectral experiments. A.Z conducted and A.Z. and M.J.M. designed and analyzed the DIA-MS proteomics experiments. A.Z. and K.Y. performed the cloning and established the FABP1 proteoform expression and purification protocol. Y.W.W. and A.Z. wrote the manuscript, which was revised by N.I. with input from all coauthors.

## Competing interests

N.I. has served as a paid consultant for Merck & Co Inc and for Boehringer Ingelheim Ltd during the past three years. She receives research funding from Amgen and Pfizer and has received honoraria from Pfizer, Elsevier and National Institutes of Health during the past three years all unrelated to this publication. The MacCoss Lab at the University of Washington has a sponsored research agreement with Thermo Fisher Scientific, the manufacturer of the instrumentation used in this research. M.J.M. is a paid consultant for Thermo Fisher Scientific. K.C.Y. is currently employee of Bristol Myers Squibb. The other authors have no competing interests.

**Correspondence** and requests for materials should be addressed to Nina Isoherranen

