## Supplemental results and materials for "Liver Fatty Acid Binding Protein (FABP1) Affects Cytochrome P450 2C9 (CYP2C9)-Mediated Metabolism in Human Liver Models"

#### Supplementary Results

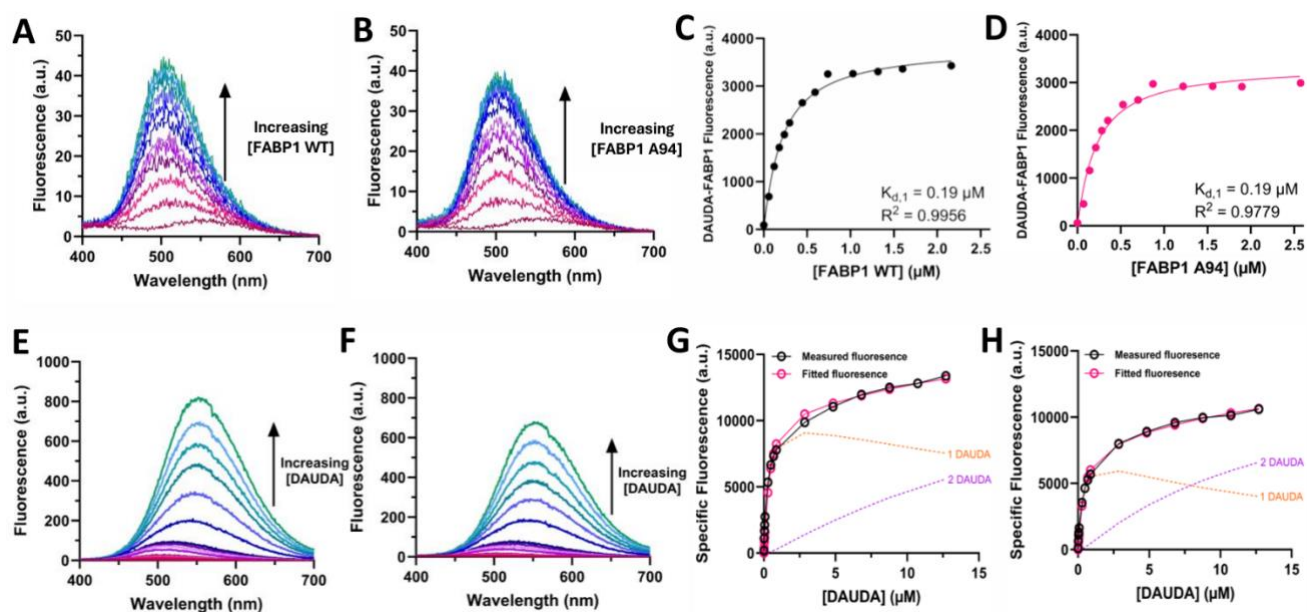

**Supplementary Fig. 1. Characterization of DAUDA binding to FABP1 WT and A94 by fluorescence spectroscopy.** **A-B** Representative raw spectra of reverse titration, where FABP1 WT or A94 was titrated in a fixed concentration of DAUDA. **C-D** Non-linear fits of DAUDA-FABP1 fluorescence from reverse titration with FABP1 WT and A94. **E-F** Representative raw spectra of forward titration, where DAUDA was titrated in a fixed concentration of FABP1 WT or A94. **G-H** Specific fluorescence from forward titrations with FABP1 WT or A94. Black circles indicate total fluorescence from DAUDA-FABP1 complexes and pink circles indicate the fitted value (DAUDA-FABP1[orange dashed line] + DAUDA-FABP1-DAUDA[purple dashed line]) at each data point in sample WT (left) and A94 (right) forward titrations, with fluorescence of free DAUDA removed by SVD deconvolution.<sup>1</sup>

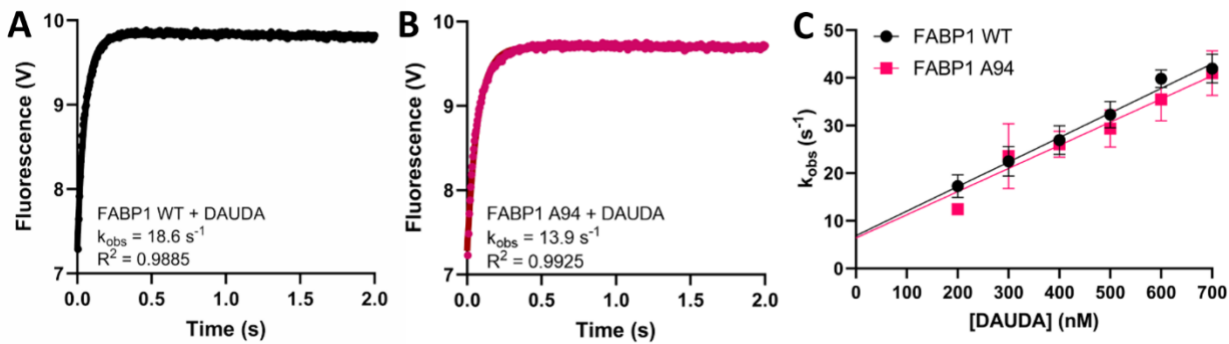

**Supplementary Fig. 2. Characterization of DAUDA binding kinetics to FABP1 WT and A94 using stopped-flow spectroscopy.** A-B Representative fluorescence over time plots with one-phase association fit for DAUDA. C Linear regression of $k_{obs} \text{ (s}^{-1}\text{)}$ over ligand concentrations for FABP1 WT and A94.

**Supplementary Table 1. Kinetic parameters of DAUDA binding to FABP1 determined by stopped-flow.** The 95% confidence intervals for the estimated parameters are shown in the parentheses.

| Ligand | Protein | $k_{on}$<br>($\text{nM}^{-1}\text{s}^{-1}$) | $k_{off}$<br>($\text{s}^{-1}$) | $K_d$ (nM) |
| --- | --- | --- | --- | --- |
| DAUDA | FABP1 WT | 0.05<br>(0.04-0.06) | 7.5<br>(2.9-12.2) | 150<br>(57-260) |
|  | FABP1 A94 | 0.05<br>(0.04-0.06) | 7.0<br>(3.2-10.2) | 140<br>(69-220) |

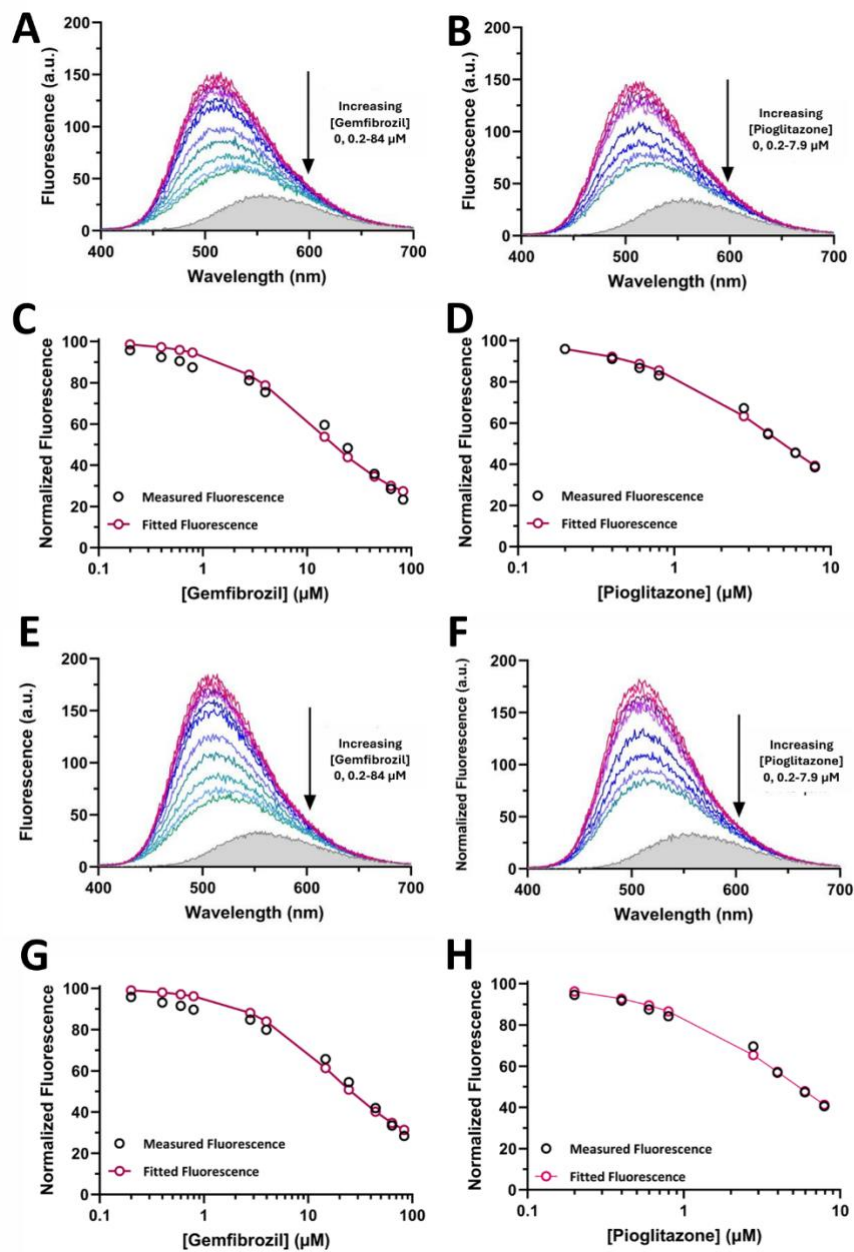

**Supplementary Fig. 3. DAUDA displacement from WT or A94 FABP1 by gemfibrozil and pioglitazone.** **A-B** The raw fluorescence spectra for DAUDA displacement titrations with WT protein and gemfibrozil (**A**) or pioglitazone (**B**) as ligand. **C-D** The corresponding DAUDA displacement curves with WT FABP1 for gemfibrozil (**C**) or pioglitazone (**D**). **E-F** The raw fluorescence spectra for DAUDA displacement titrations with A94 variant and gemfibrozil (**E**) or pioglitazone (**F**) as ligand. **G-H** The corresponding DAUDA displacement curves with A94 variant FABP1 for gemfibrozil or pioglitazone. The concentration ranges of drug ligands titrated are indicated next to the raw fluorescence spectra. The gray shaded areas show the spectrum of 0.5 $\mu\text{M}$ unbound DAUDA in the absence of FABP1 and drug ligand. Solid lines on the panels on the right indicate fits to a ternary complex binding model as previously established.<sup>1</sup> The resulting $K_d$ values are summarized in **Table 1** in the main manuscript.

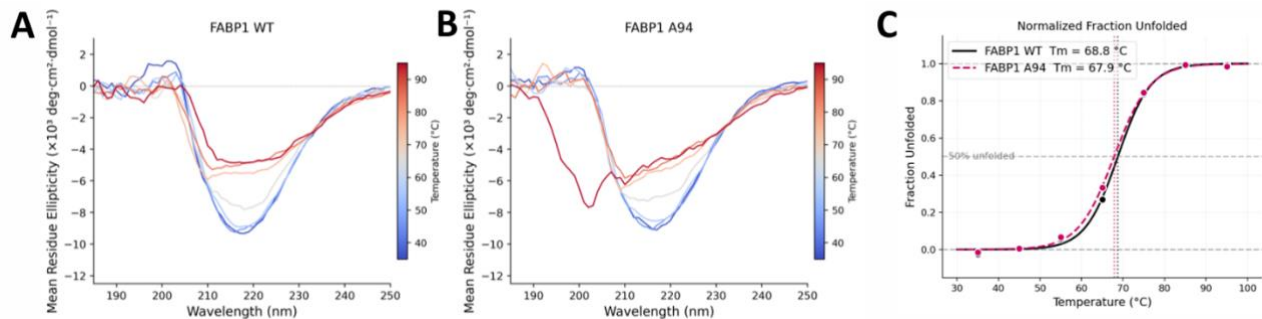

**Supplementary Fig. 4. Thermal denaturation of FABP1 WT and A94 monitored by circular dichroism.** **A-B** Far-UV CD spectra (185–250 nm) of wild-type FABP1 (**A**) and the A94 variant (**B**). Spectra are colored by temperature according to the scale bar at right (blue, low; red, high). (**C**) Fraction unfolded as a function of temperature, derived from the ellipticity at 222 nm and normalized between the folded and unfolded baselines. Solid black line and dashed magenta line in panel **C** are the fits to a two-state unfolding model for WT and A94 respectively, and symbols are experimental points. Vertical dotted lines mark the fitted melting temperatures (WT, 68.8 °C; A94, 67.9 °C); the horizontal dashed line indicates 50% unfolded.

**Supplementary Table 2. University of Washington human liver bank donor characteristics.**
Data are reported for all 61 donors except BMI, which was available for 34.

| **UW Liver Bank donor characteristics** | **N or Mean (range)** |
| --- | --- |
| Sex | Female (n = 28)<br>Male (n = 33) |
| Age (years) | 41 (7 – 68) |
| BMI (kg/m<sup>2</sup>) | 25.2 (17.0 – 44.8) |
| Race | Asian (n = 1)<br>Black (n = 3)<br>Caucasian (n = 57) |
| Liver Pathology | Normal (n = 26)<br>Fatty (n = 26)<br>[Fatty (n=17), Mildly Fatty (n=7), and Moderate<br>Microvesicular Fat (n=2)]<br>Acute Injury (n=4)<br>Fibrotic (n=4)<br>Iron Deposition (n=1) |

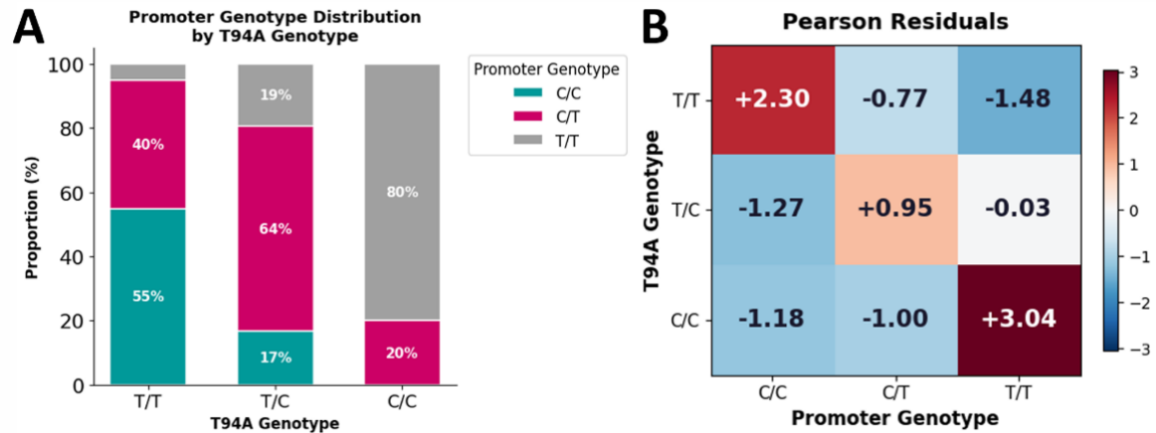

**Supplementary Fig. 5. Association between the FABP1 promoter genotype and the FABP1 T94A genotype.** **A** Distribution of FABP1 promoter genotype stratified by T94A genotype, expressed as the proportion (%) of samples within each T94A group. **B** Pearson residuals from a chi-square test of independence between promoter genotype (columns) and T94A genotype (rows).

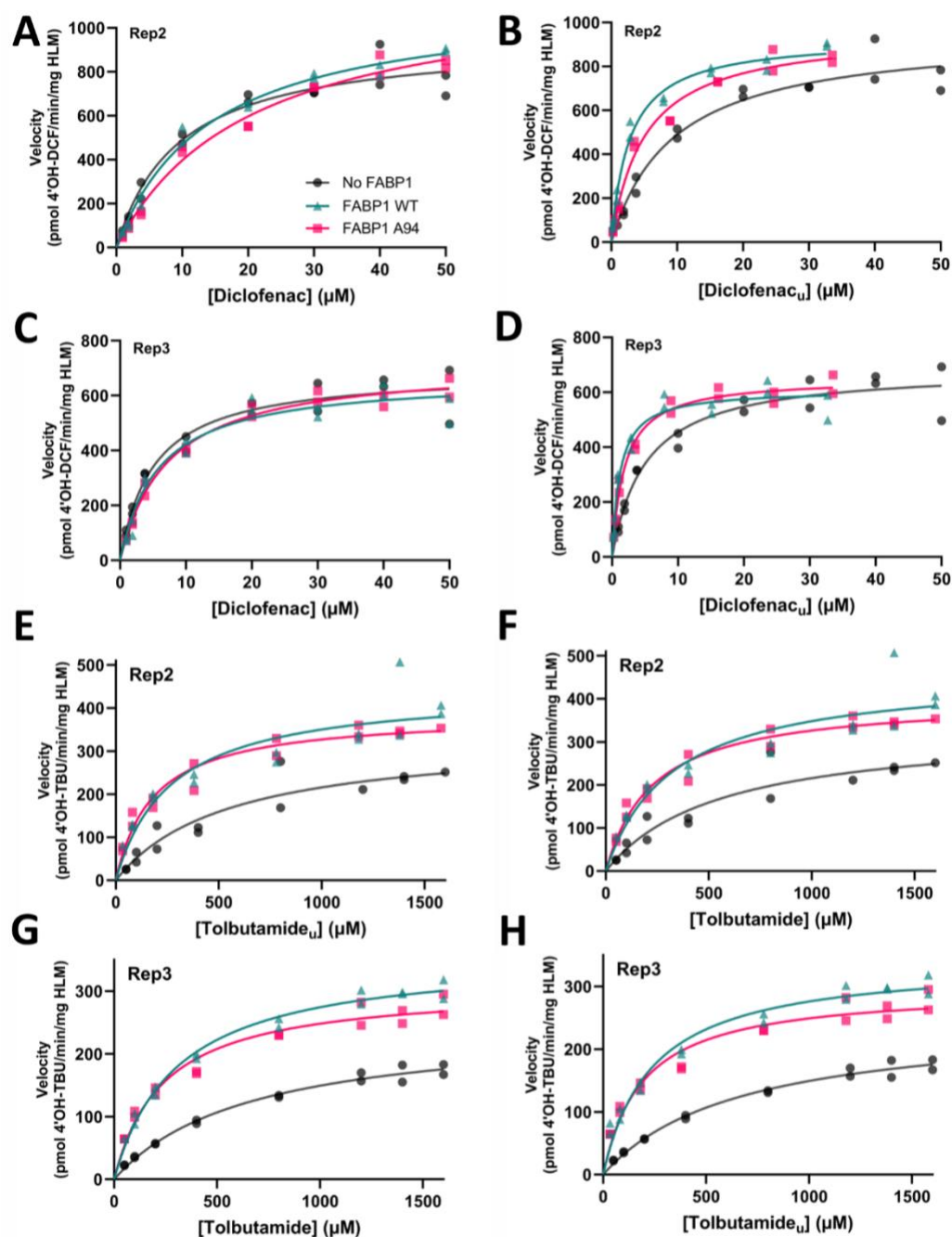

**Supplementary Fig. 6. Additional replicates supporting main manuscript Figure 4 showing the effect of FABP1 on CYP2C9-mediated metabolism of diclofenac and tolbutamide in pooled human liver microsomes (HLM) with or without 20 mM FABP1 WT or A94 variant.** The formation rate of 4'-hydroxydiclofenac (4'OH-DCF) as a function of total diclofenac concentration (A,C) and as a function of unbound diclofenac concentration (B,D). The formation rate of 4'-hydroxytolbutamide (4'OH-TBU) as a function of total tolbutamide concentration (E,G) and as a function of unbound tolbutamide concentration (F,H). The points are individual measured velocities within the experiment, and solid lines are nonlinear least-squares fits of the Michaelis–Menten equation to the data. The kinetic parameters are summarized in main manuscript **Table 2**.

**Supplementary Table 3. Michaelis–Menten kinetic parameters for CYP2C9-mediated metabolism of total diclofenac and tolbutamide in human liver microsomes in the absence or presence of 20 mM FABP1 WT or A94 variants.** For each substrate, parameters are reported relative to total (nominal) substrate concentrations. Values are shown as mean ± standard deviation of three independent experiments with all three groups done together. Each parameter was compared between no FABP1 and FABP1 WT, no FABP1 and FABP1 A94, and FABP1 WT and FABP1 A94 using ratio paired t-tests for each pairwise comparison.

|  | **Diclofenac** |  |  | **Tolbutamide** |  |  |
| --- | --- | --- | --- | --- | --- | --- |
|  | No FABP1 | +FABP1<br>WT | +FABP1 A94 | No FABP1 | +FABP1<br>WT | +FABP1 A94 |
| $V_{\max}$<br>(pmol/min/mg) | 768 ± 160 | 852 ± 248<sup>n.s.</sup> | 885 ± 277<sup>n.s.,n.s.</sup> | 276 ± 50 | 389 ± 61<sup>**</sup> | 364 ± 49<sup>n.s.,n.s.</sup> |
| $K_m$<br>(μM) | 7.4 ± 2.2 | 9.3 ± 4.4<sup>n.s.</sup> | 12.3 ± 7.1<sup>n.s.,n.s.</sup> | 621 ± 61 | 307 ± 13<sup>*</sup> | 250 ± 28<sup>*, n.s.</sup> |
| $CL_{int}$<br>(μL/min/mg) | 107 ± 25 | 97 ± 18<sup>n.s.</sup> | 79 ± 20<sup>n.s.,n.s.</sup> | 0.45 ± 0.13 | 1.26 ± 0.14<sup>*</sup> | 1.46 ± 0.25<sup>**,*</sup> |

The raw $p$ values were corrected for multiple comparisons by the Holm–Bonferroni procedure applied separately within each kinetic parameter, treating the three pairwise comparisons of a given parameter (no FABP1 vs. FABP1 WT, no FABP1 vs. FABP1 A94, and FABP1 WT vs. FABP1 A94) as one family of tests. Significance is reported at a family-wise error rate of $\alpha = 0.05$. The superscript of “+ FABP1 WT” indicates the significance for the no FABP1 vs. FABP1 WT comparison. The superscripts of “+ FABP1 A94” indicate the significance for no FABP1 vs. FABP1 A94 comparison (first) and FABP1 WT vs. FABP1 A94 comparison (second). n.s., not significant; \*, adjusted $p$ value < 0.05; \*\*, adjusted $p$ value < 0.01.

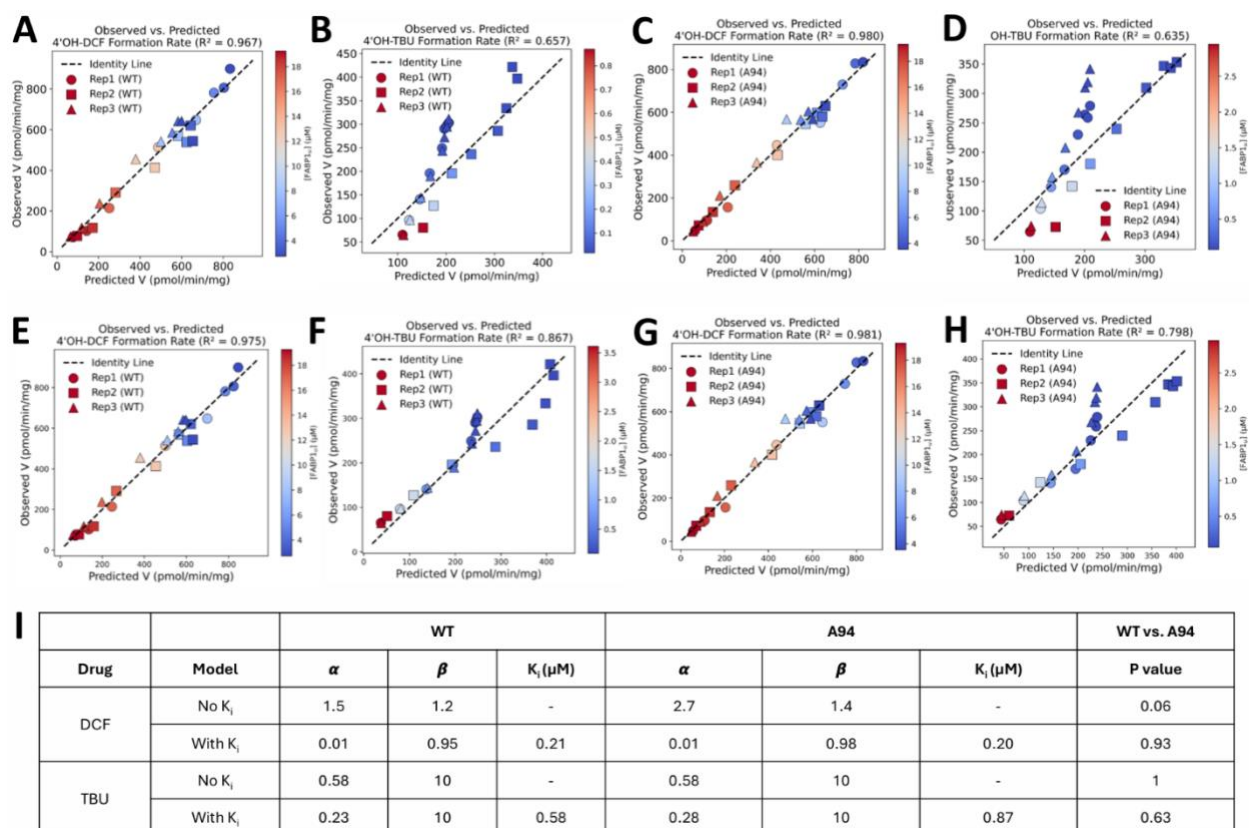

**Supplementary Fig. 7. Estimation of kinetic parameters for 4'OH-diclofenac (4'OH-DCF) and 4'OH-tolbutamide (4'OH-TBU) with FABP1 WT and A94.** A-B Observed versus predicted 4'OH-DCF or 4'OH-TBU formation rate assuming no apo-FABP1-CYP2C9 interaction model (no $K_i$ model) for WT FABP1. C-D Observed versus predicted 4'OH-DCF or 4'OH-TBU formation rate assuming no $K_i$ model for A94 FABP1. E-F Observed versus predicted 4'OH-DCF and 4'OH-TBU formation rate assuming with apo-FABP1-CYP2C9 interaction model (with $K_i$ model) for WT FABP1. G-H Observed versus predicted 4'OH-DCF and 4'OH-TBU formation rate assuming with $K_i$ model for A94 FABP1. I Summary table of best fitted $\alpha$, $\beta$, and $K_i$ assuming models in panels A-H for diclofenac and tolbutamide respectively. No $K_i$ and with $K_i$ models were compared using small-sample-corrected Akaike Information Criterion (AICc) and Bayesian Information Criterion (BIC). The fitted parameters of each model were compared globally using extra sum-of-squares F-test for diclofenac and tolbutamide respectively.

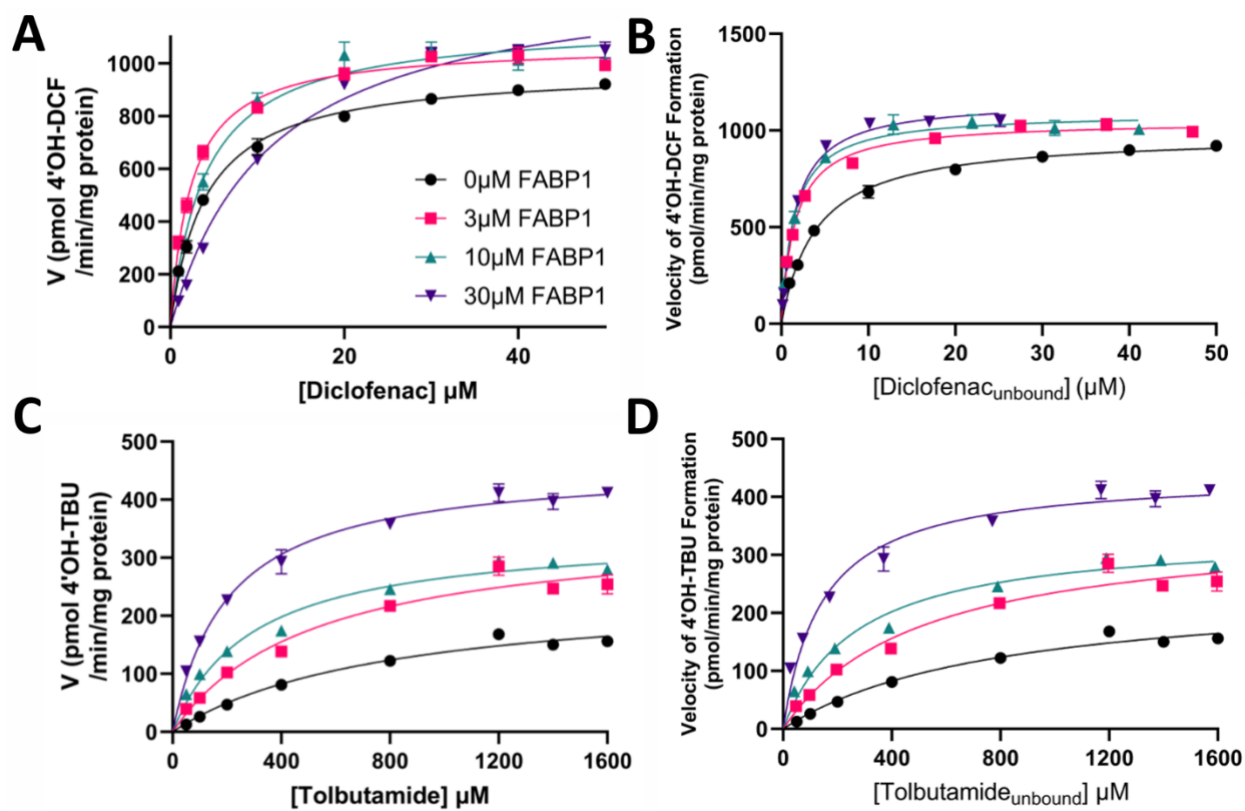

**Supplementary Fig. 8. Effect of FABP1 on CYP2C9-mediated metabolism of diclofenac and tolbutamide in human liver microsomes (HLM) with or without varying concentration of FABP1 WT.** **A** Formation rate of 4'-hydroxydiclofenac (4'OH-DCF) as a function of total diclofenac concentration. **B** Formation rate of 4'OH-DCF as a function of unbound diclofenac concentration. **C** Formation rate of 4'-hydroxytolbutamide (4'OH-TBU) as a function of total tolbutamide concentration. **D** Formation rate of 4'OH-TBU as a function of unbound tolbutamide concentration. In all panels, points are average measured velocities with range indicated as error bars, and solid lines are nonlinear least-squares fits of the Michaelis–Menten equation to the data.

**Supplementary Table 4. Comparison of linear regression models predicting CYP2C9-mediated metabolite formation from CYP2C9 relative abundance and FABP1 expression levels.** Model notation follows the form outcome ~ predictor(s). For ln(4'OH-diclofenac formation rate) and ln(4'OH-tolbutamide formation rate), a base model containing only ln(CYP2C9) was compared with a series of models adding total FABP1 or FABP1 WT + FABP1 A94. Goodness of fit is summarized by the coefficient of determination ($R^2$), adjusted $R^2$, the small-sample-corrected Akaike information criterion (AICc), and the Bayesian information criterion (BIC). P values were obtained from F tests comparing each model with its respective base model (nested-model comparison). The base model is the reference and therefore has no associated p value. Raw p values are shown alongside p values adjusted for multiple comparisons using the Holm step-down procedure. 4'OH-DCF, 4-hydroxydiclofenac; 4'OH-TBU, 4'hydroxytolbutamide.

| Model | $R^2$ | Adj.<br>$R^2$ | AICc | BIC | Raw p<br>values | Adj. p<br>values |
| --- | --- | --- | --- | --- | --- | --- |
| Base: ln(4'OH-DCF) ~ ln(CYP2C9) | 0.479 | 0.470 | 137 | 141 | - | - |
| ln(4'OH-DCF) ~ ln(CYP2C9)+FABP1 | 0.544 | 0.528 | 131 | 137 | 0.0058 | 0.012 |
| ln(4'OH-DCF) ~ ln(CYP2C9)+FABP1 WT + FABP1 A94 | 0.547 | 0.523 | 133 | 141 | 0.019 | 0.019 |
| Base: ln(4'OH-TBU) ~ ln(CYP2C9) | 0.508 | 0.500 | 133 | 137 | - | - |
| ln(4'OH-TBU) ~ ln(CYP2C9)+FABP1 | 0.567 | 0.552 | 127 | 133 | 0.0068 | 0.014 |
| ln(4'OH-TBU) ~ ln(CYP2C9)+FABP1 WT + FABP1 A94 | 0.569 | 0.547 | 129 | 137 | 0.023 | 0.023 |

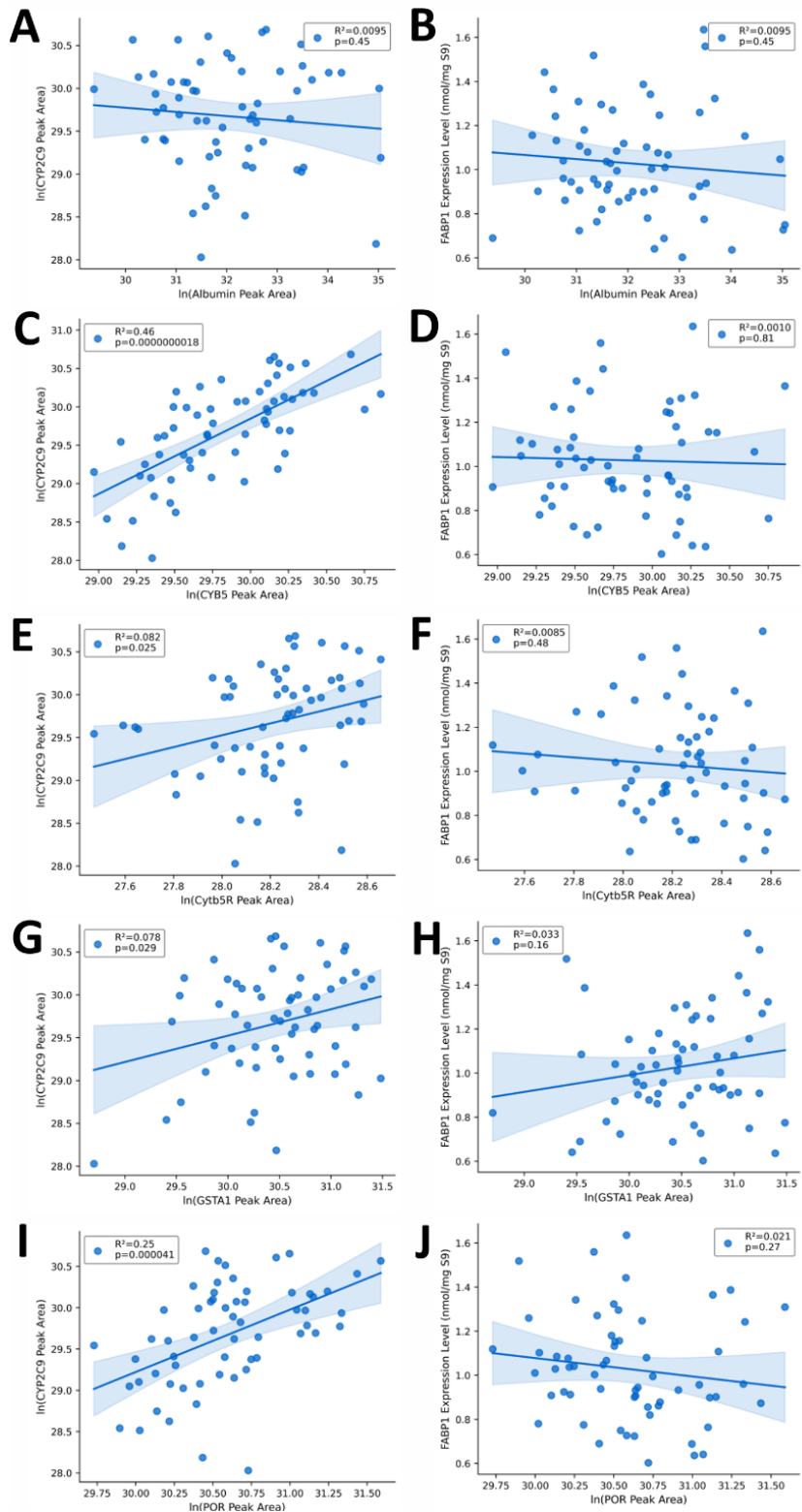

**Supplementary Fig. 9. Correlations between CYP2C9 relative abundance or FABP1 expression level with a list of proteins that potentially affect CYP2C9-mediated metabolisms.** **A** Scatter plot of ln(CYP2C9 peak area) versus ln(Albumin peak area). **B** Scatter plot of FABP1 expression level versus ln(Albumin peak area). **C** Scatter plot of ln(CYP2C9 peak area) versus ln(CYB5 peak area). **D** Scatter plot of FABP1 expression level versus ln(CYB5 peak area). **E** Scatter plot of ln(CYP2C9 peak area) versus ln(Cytb5R peak area). **F** Scatter plot of FABP1 expression level versus ln(Cytb5R peak area). **G** Scatter plot of ln(CYP2C9 peak area) versus ln(GSTA1 peak area). **H** Scatter plot of FABP1 expression level versus ln(GSTA1 peak area). **I** Scatter plot of ln(CYP2C9 peak area) versus ln(POR peak area). **J** Scatter plot of FABP1 expression level versus ln(POR peak area). Scatter plots with best fitted linear regression line and 95% confidence intervals were shown for each panel. Blue circles show the individual data point in 61 HLS9s. CYP2C9, Cytochrome P450 2C9; CYB5, cytochrome b5; Cytb5R, NADH cytochrome b5 reductase; GSTA1, glutathione S-transferase A1; POR, cytochrome P450 oxidoreductase; FABP1, FABP1. The coefficient of determination ($R^2$) and p values acquired from Pearson's correlation analysis are shown in the legend on each panel.

### Supplementary Methods

#### I. Chemicals and Reagents

Kanamycin, Trizma base (Tris), sodium chloride, sodium phosphate, potassium phosphate, potassium chloride, benzonase, butanol, thrombin, Rosetta 2 *E. coli*, Coomassie Brilliant Blue R, 11-(Dansylamino)-undecanoic acid (DAUDA), β-Nicotinamide-adenine dinucleotide phosphate (NADPH) tetrasodium salt, iodoacetamide (IAC), EDTA-free protease inhibitor cocktail, tolbutamide, and 4-hydroxytolbutamide were purchased from Millipore-Sigma (St. Louis, MO). 4-hydroxydiclofenac and 4'-OH-5-chloro-diclofenac were a gift from Dr. Allan Rettie (Department of Medicinal Chemistry, University of Washington). Tryptone, yeast extract, isopropyl β-D-thiogalactopyranoside (IPTG), phenylmethyl sulfonyl fluoride (PMSF), dithiothreitol (DTT), imidazole, sodium dodecyl sulfate (SDS), Pierce bicinchoninic acid (BCA) protein assay, high-performance liquid chromatography (HPLC) and mass spectrometry (Optima) grade acetonitrile, methanol, water and formic acid were purchased from Thermo Fisher Scientific (Waltham, MA). Lipidex-5000 suspension was purchased from Perkin Elmer Inc (Waltham, MA, USA). Glycine and Mini-PROTEAN TGX protein gels were purchased from Bio-Rad (Hercules, CA). Mass spectrometry grade recombinant trypsin was ordered from Promega (Madison, WI). Stable isotope-labeled peptide (SIL-peptide) labeled with phenylalanine (+10 Da) ($F[^{13}C_9,^{15}N]$TITAGSK) was ordered from Thermo Fisher Scientific (Waltham, MA) and used as internal standard for FABP1 quantification.

#### II. Human FABP1 A94 Cloning

FABP1 cDNA was obtained from OriGene Technologies (Rockville, MD) (catalog number SC119222). Primers were designed (forward primer sequence: 5'- caccatgagtttctccggcaagtac-3'; reverse primer sequence: 5'- aattctcttgctgattctcttgaag-3') to amplify the FABP1 coding sequence and allow cloning into Novagen's pET28a Expression vector according to the manufacturer's instructions. This system adds an N-terminal 6xHis tag to the amplified coding sequence to allow *E. coli* expression and downstream purification. The resulting plasmid was sequenced and confirmed to be correct, and to contain the previously described single nucleotide polymorphism (SNP) resulting in the A94 variant of this protein.<sup>2</sup>

**A**

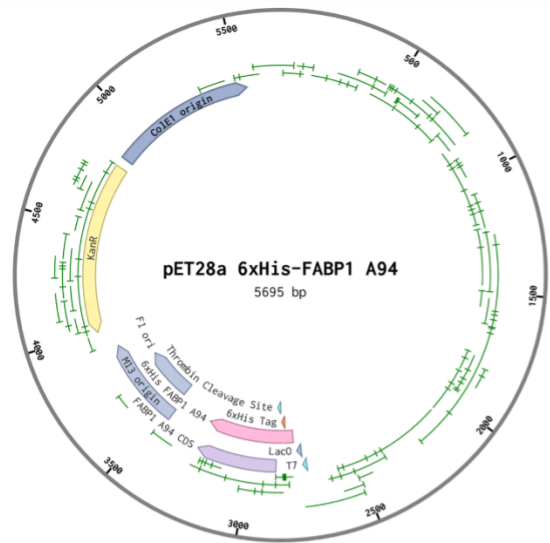

**B**

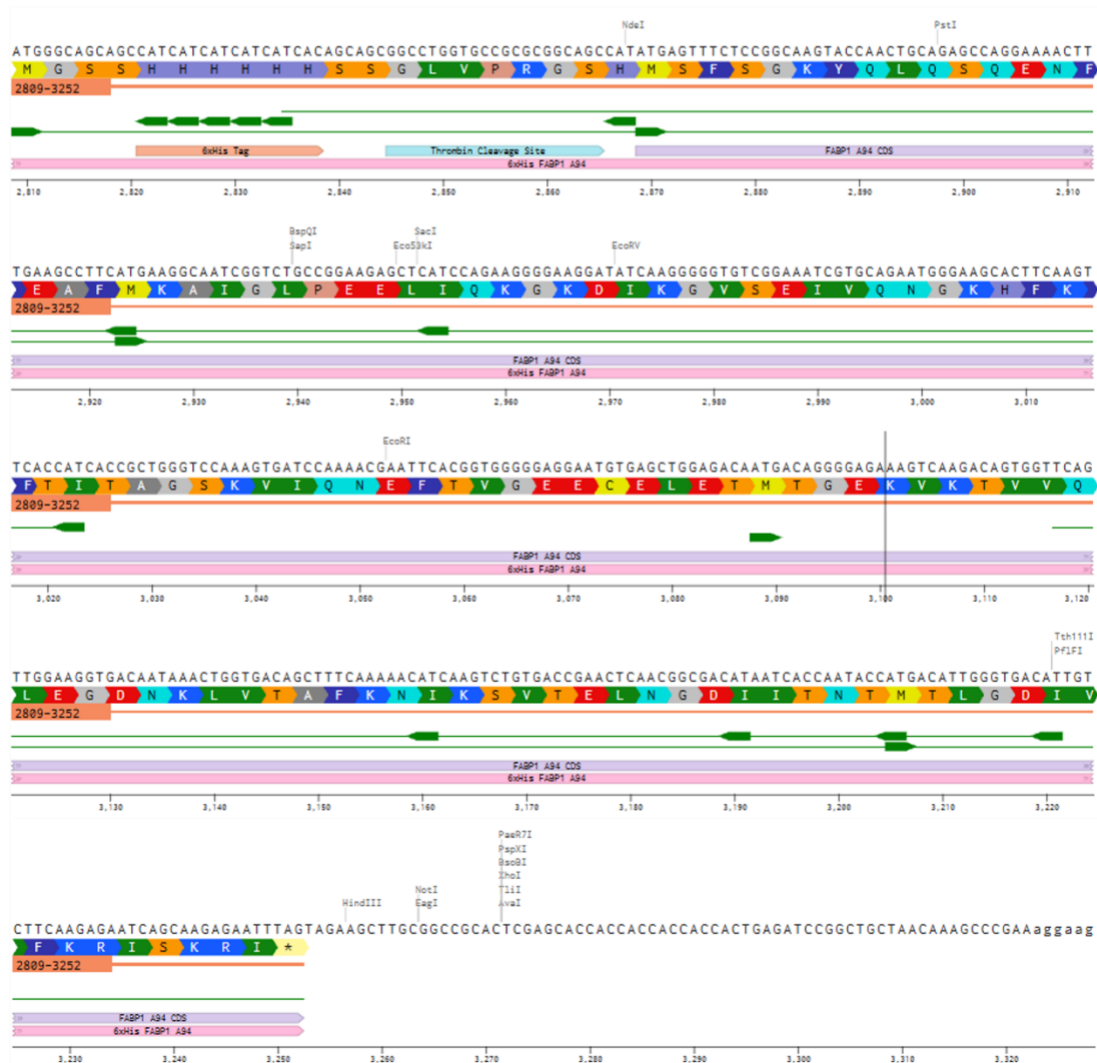

**Supplementary Fig. 10. Plasmid map and coding sequence for recombinant FABP1 A94 visualized by Benchling (2024). Retrieved from https://benchling.com.**

##### III. Expression and purification of human FABP1 WT and A94 variants

FABP1 WT and A94 variants were expressed and purified using the same protocol as described in this section. Both 6xHis-tagged WT and A94 mutant human FABP1 were expressed in Rosetta 2 *E. coli* (Novagen, Madison, WI) and were purified using a DuoFlow fast protein liquid chromatograph (FPLC) (Bio-Rad, Hercules, CA) as previously described<sup>3</sup> with minor modifications. Briefly, his-tagged FABP1 was purified using a 1 mL HisTrap HP affinity column (GE Healthcare, Chicago, IL) with 20 mM Tris, pH 7.4, 500 mM NaCl, 30 mM imidazole as the binding buffer and 20 mM Tris, pH 7.4, 500 mM NaCl, 350 mM imidazole for elution. The his-tag was cleaved by incubation with thrombin (0.01 U thrombin per 10 mg FABP1) for 16-hr on ice in 20mM Tris, pH 7.4, 300 mM NaCl and 200 mM imidazole. Cleaved FABP1 was buffer exchanged using a Superdex 75 (Marlborough, MA) column into 50 mM potassium phosphate buffer pH 7.4, 100 mM potassium chloride. Delipidation of the cleaved FABP1 was performed using three rounds of butanol treatment (5 mL butanol to 5 mL cleaved FABP1) and 30-minute incubation with preconditioned Lipidex-5000 (Perkin Elmer Inc., Waltham, MA, USA; 0.1g per 1 mg of FABP1) stored in 50 mM potassium phosphate pH 7.4, 100 mM KCl as previously described<sup>3</sup>. The concentration of purified FABP1 was quantified via bicinchoninic acid (BCA) assay (Pierce, Waltham, MA) and the quality of the protein was confirmed by fluorescence titration with DAUDA. The delipidated FABP1 was flash frozen with addition of 5% glycerol and stored at -80°C for fluorescent binding, protein quantification, and enzymatic activity assays.

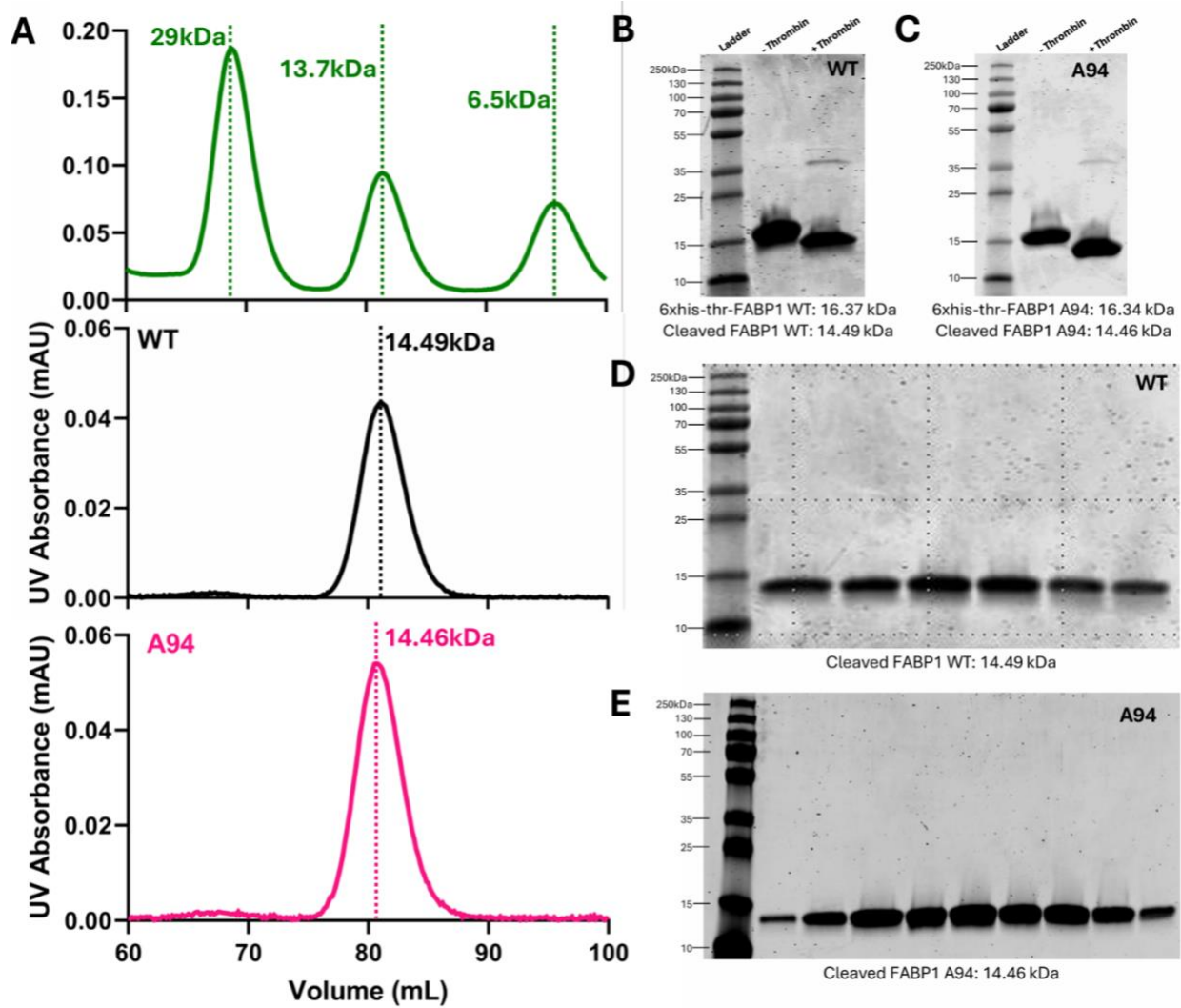

**Supplementary Fig. 11. Purification of recombinant FABP1 WT and A94.** **A** Size-exclusion chromatography of protein standards, cleaved FABP1 WT, and cleaved FABP1 A94 generated with Superdex 75 size exclusion column (GE Healthcare, Chicago, IL). Both variants elute as a single symmetric peak **B-C** Coomassie-stained SDS-PAGE (anyKD gel) of WT and A94 before (–Thrombin) and after (+Thrombin) cleavage of the N-terminal 6×His-thrombin tag. **D-E** Coomassie-stained SDS-PAGE of pooled peak fractions from the size-exclusion runs in panel **A** for WT and A94, showing a single band at the expected mass with no detectable contaminants. Molecular weight markers (kDa) are indicated at left of each gel.

#### IV. Characterization of FABP1 WT and A94 variants

##### 1. Estimation of DAUDA binding affinity to FABP1 WT and A94 using fluorescence spectroscopy

All fluorescence spectra were collected using a Cary Eclipse Fluorescence Spectrophotometer (Agilent, Santa Clara, CA) using 4 mL clear quartz cuvettes. The fluorescence assays were adapted from previously established methods.<sup>1</sup> Briefly, fluorescence spectroscopy experiments were done in 2 mL of assay buffer (50 mM potassium phosphate, pH 7.4, 100 mM KCl) at 21°C. The final concentration of organic solvent was kept at < 2%. All experiments were repeated on at least three separate days and with at least two independent batches of purified protein.

The equilibrium binding affinity ($K_d$) for DAUDA with FABP1 was determined using reverse and forward fluorescence titrations as previously established.<sup>1</sup> Briefly, in the reverse titrations, FABP1 (0.04 – 2.6 μM) was added to 0.05 μM DAUDA. In forward titrations DAUDA (0.02 – 12.7 μM) was added to FABP1 (0.3 μM). The emission spectra of fluorescent probe and probe-FABP1 overlapped preventing direct measurement of probe-FABP1 fluorescence in titration experiments. Singular value decomposition (SVD) was therefore used to deconvolute fluorescence titration spectra as previously described.<sup>1</sup> The high affinity equilibrium dissociation constant ($K_{d,1}$) was estimated by fitting a single-site binding model to the reverse titration data. The binding affinity for a low binding site ($K_{d,2}$) was estimated by fitting a sequential two-site binding model to the forward titration data using fixed $K_{d,1}$ from the reverse titration fit as previously reported.<sup>1</sup> Model fitting was done with COPASI as previously published.<sup>1</sup> $K_d$ values are reported (± standard deviation) from independent replicate data sets generated on different days with different batches of purified FABP1 WT or A94.

##### 2. Estimation of DAUDA binding kinetics to FABP1 WT and A94 using stopped flow

Kinetics of DAUDA binding to wild-type and A94 variant FABP1 was measured using an SX20 stopped-flow instrument (Applied Photophysics, Leatherhead, Surrey, UK) with a 1 ms dead time. A 2 mm optical path length was used, and the detector voltage was autoPMed (automatic photomultiplier) for each ligand concentration (range: 300 - 450 volts). To measure the kinetics of DAUDA binding to FABP1 the excitation wavelength was set to 320 nm and the increase in FABP1-DAUDA fluorescence was monitored using a 365 nm emission cut-off filter. All binding experiments were carried out at room temperature.

To measure DAUDA binding kinetics with FABP1, separate solutions of 400 nM FABP1 WT or 400 nM FABP1 A94 and varying concentrations of DAUDA (400, 600, 800, 1000, 1200, 1400 nM) were prepared in an assay buffer (50 mM potassium phosphate pH 7.4, 100 mM KCl) in clear glass vials. Each ligand solution contained 1% methanol. Protein and ligand solutions were loaded into separate 1 mL drive syringes. After injection, the final mixed concentrations were 200 nM FABP1 and 12.5, 100, 200, 300, 400, 500, 600 or 700 nM for DAUDA. Data were acquired for 2 seconds with 400 measurements per injection for DAUDA. At least two injections were conducted with each ligand concentration, and three replicate experiments were conducted on three different days for both FABP1 variants and ligand combinations.

To determine the kinetic parameters of DAUDA to FABP1 WT or A94, a one phase association exponential model was used to estimate observed association constant ($k_{obs}$) for each ligand concentration in GraphPad Prism (V 10.1.2). A simple linear regression model against $k_{obs}$ and ligand concentration was used to acquire slope ($k_{on}$) and y-intercept ($k_{off}$) in GraphPad Prism (V 10.1.2). $K_d$ was estimated using the ratio between $k_{off}$ and $k_{on}$ for DAUDA and FABP1 binding.

##### 3. Estimation of thermal stability of FABP1 WT and A94

Thermal stability of FABP1 WT and A94 was determined using JASCO J1500 Circular Dichroism spectrometer (JASCO, Easton, MD) equipped with a Koolance EXOS liquid cooling system. All experiments (temperature and ligand interaction) were done at a final protein concentration of 0.5 μM in a buffer containing 10 mM potassium phosphate (pH 7.4). Each protein sample was incubated at 25 °C for 10 min prior to scanning. The sample was scanned 10 times from 185 nm to 250 nm. The final CD spectrum obtained represented an average of ten scans, the spectrum was background-subtracted, and the spectrum was mathematically smoothed using the Means-Movement method with a convolution width = 5. Thermal unfolding was monitored by extracting the signal at 222 nm which is the wavelength that is most sensitive to α-helical content at each measured temperature. The resulting melt curve for each protein sample was fitted to a two-state unfolding model in which the observed signal varies sigmoidally with temperature.

**Supplementary Table 5. Mass spectrometer multiple reaction monitoring (MRM) transitions and parameters for human specific and unique tryptic peptides of FABP1 WT and A94 variants.**

| Protein | Peptide | Precursor Ion ($m/z$) | Fragment Ion ($m/z$) | DP (V) | CE (V) |
| --- | --- | --- | --- | --- | --- |
| FABP1 WT | LVTTFK<br>(+2) | 355 | y4<sup>+</sup> 496 | 80 | 16 |
|  |  |  | y5<sup>+</sup> 595 | 80 | 16 |
|  |  |  | y3<sup>+</sup> 395 | 80 | 16 |
| FABP1 A94 | LVTAFK<br>(+2) | 340 | y4<sup>+</sup> 466 | 80 | 16 |
|  |  |  | y3<sup>+</sup> 365 | 80 | 16 |
|  |  |  | y5<sup>+</sup> 565 | 80 | 16 |

#### V. LC-MS/MS quantification of 4'OH-diclofenac and 4'OH-tolbutamide

4'OH-5-chloro-diclofenac was used as the internal standard for 4'OH-DCF quantification whereas 4'OH-mephenytoin-d<sub>3</sub> (4'OH-MPH-d<sub>3</sub>) was used as the internal standard for 4'OH-TBU quantification. 4'OH-DCF and 4'OH-TBU were separated using a Zorbax Eclipse Plus C18 column (2.1 x 50 mm, 5 µM, Agilent Technologies, Santa Clara, CA) with the column oven set to 35 °C. Mobile phases A and B consisted of water and acetonitrile, respectively, both containing 0.1 % formic acid. A gradient elution at a flow rate of 0.4 mL/min was used as follows: mobile phase A was kept at 95% and B at 5% for the first 0.1 min, then B was increased to 70% and 95% by 3.5 and 4.0 min respectively, returned to initial conditions by 5 min, and the column was re-equilibrated in initial conditions for an additional 1.5 min. All analytes were monitored in positive ion mode with electrospray ionization and the MS parameters were set as follows: IS 5000 V, TEM 600 °C, CUR 35 p.s.i., GS1 50, GS2 50, CAD medium, DP 90 and 1 (4'OH-DCF and 4'OH-5-chloro-diclofenac), EP 10 and 10 (4'OH-DCF and 4'OH-5-chloro-diclofenac, respectively), CE 30 and 19 (4'OH-DCF and 4'OH-5-chloro-diclofenac, respectively), CXP 15 and 16 V (4'OH-DCF and 4'OH-5-chloro-diclofenac, respectively). Two different MRM transitions (312>230 *m/z*, and 346>300 *m/z*) were used to monitor 4'OH-DCF and 4'OH-5-chloro-diclofenac, respectively. Quantification of 4'OH-TBU was done using the same analytical method published from us previously.<sup>4</sup> Signal-to-noise (S/N) cutoffs of 3 and 5 were used as the lower limit of detection (LLOD) and lower limit of quantification (LLOQ). The linearity of detector response for all the metabolites was verified. All analytes were quantified based on a standard curve ranging from 2.34 nM to 300 nM for 4'OH-TBU and 4'OH-DCF quantification, and from 1.56 to 200 nM for 4'OH-DCF quantification.

#### VI. Kinetics of 4'-OH-diclofenac and 4'-OH-tolbutamide formation with varying concentrations of FABP1 in pooled HLMs

The kinetics of 4'OH-DCF and 4'OH-TBU formation in the presence and absence of FABP1 in pooled HLMs were determined under conditions of protein and time linearity. HLMs pooled from 50 donors were purchased from Gibco (Cat. HMMCPL, Thermo Fisher Scientific, Waltham, MA). Pooled HLMs (0.01 mg/mL protein) with or without 3, 10, or 30 µM WT FABP1 were preincubated with eight different concentrations of diclofenac (0.9 – 50 µM) or tolbutamide (20 – 1600 µM) for 5 minutes at 37 °C in 90 µL incubation buffer (10 mM potassium phosphate, pH 7.4, 150 mM potassium chloride, 0.5 mM DTT) in a 96-well plate. Reactions were initiated with 1 mM NADPH (final concentration) to a final volume of 100 µL and quenched after 10 minutes by 100 µL of the ice-cold acetonitrile containing 25 nM 4'OH-5-chloro-diclofenac or 10 nM 4'OH-mephenytoin as an internal standard for diclofenac or tolbutamide assay separately. Experiments with and without FABP1 were done matched on the same day with technical duplicates. The quenched reactions were centrifuged at 3,200 g for 60 minutes at 4 °C and 100 µL supernatant was collected and transferred to 96-well MS plate for LC-MS/MS analysis.

#### VII. Global Proteomics

##### 1. Protein Digestion

Human liver S9 fractions were prepared for mass spectrometry analysis using protein aggregation capture (PAC) as described previously<sup>5</sup> with minor modifications. Individual liver S9 fractions were added to a KingFisher 96-well deep-well plate (Thermo Fisher Scientific, cat. no. 95040450) using a variable volume of liver S9 fractions plus a variable volume of diluent (50 mM potassium phosphate, 250 mM sucrose, pH 7.4) to yield 50 µg total protein in 50 µL final volume per well. The completed KingFisher plate was sealed and stored at -80°C prior to digestion. The randomized sample plate contained 61 individual liver samples and six pooled reference samples. The reference pool was generated by combining material from all 61 individual liver samples. One pooled reference sample was included in each row of the plate.

The frozen KingFisher plate was thawed at 37°C for approximately 10 minutes. Each well was brought to a final volume of 200 µL by addition of 150 µL of 100 mM Tris, pH 8.5, 2% SDS containing yeast enolase (Sigma, cat no. E6126) and tris(2-carboxyethyl)phosphine (TCEP). This resulted in final concentrations of approximately 1.5% SDS, 800 ng yeast enolase per well, and 10 mM TCEP. Enolase was included in each sample as a process control. Samples were mixed by pipetting, covered with a plate lid, vortexed at 1,200 rpm for 30 seconds in an Eppendorf ThermoMixer, and reduced for 1 hour at 37°C with shaking at 650 rpm in a ThermoMixer.

Following reduction, samples were cooled to room temperature and alkylated by addition of 10 µL of 300 mM iodoacetamide per well, yielding a final concentration of approximately 15 mM. Samples were vortexed at 1,200 rpm for 30 seconds and incubated for 30 minutes at room temperature in the dark. Alkylation was quenched by addition of 10 µL of 300 mM dithiothreitol (DTT) per well. Each well then received 12.5 µL of MagResyn Hydroxyl particles (ReSyn Biosciences, cat no. MR-HYX002), and proteins were aggregated onto the beads by addition of 767 µL acetonitrile bringing the final concentration to approximately 77%. Protein capture, washing, and digestion were performed on a KingFisher Apex (Thermofisher Scientific) as previously described.<sup>5</sup> Beads were washed in three deep-well plates containing 95% acetonitrile followed by two plates containing 70% ethanol. Protein-containing beads were released into 150 µL of 50 mM Tris, pH 8.5, containing trypsin (Pierce, cat no. PI90058) at an approximate 1:20 enzyme-to-substrate ratio. Digestion was performed in the KingFisher at 47°C for 1 hour.

Following digestion, 100 µL of each digest was transferred to a Thermo Scientific SureSTART autosampler plate (cat no. 60180-P207B) containing 7 µL of 10% trifluoroacetic acid and mixed by pipetting. A 50 µL aliquot of acidified peptides was then transferred to a second autosampler plate containing 5 µL of 500 fmol/µL Pierce Peptide Retention Time Calibration (PRTC) Mixture (Thermo Scientific, cat no. PI-88321) in 0.1% TFA. PRTCs were included as a process control.<sup>6</sup> Peptide plates were sealed and stored at -80°C until LC-MS/MS analysis.

#### 2. Mass Spectrometry

Prepared liver peptide samples were analyzed by data-independent acquisition mass spectrometry (DIA-MS). For each sample 6 $\mu\text{L}$, corresponding to 2 $\mu\text{g}$ total liver protein plus 32 ng yeast enolase and 300 fmol of PRTC, was injected onto a Thermo Fisher Scientific Orbitrap Eclipse Tribrid mass spectrometer using a “trap and elute” setup on a Thermo Vanquish Neo UHPLC System. The system was equipped with a Thermo Scientific PepMap Neo Trap Cartridge (cat no. 174500) and a Bruker PepSep C18 15 cm × 150 $\mu\text{m}$, 1.9 $\mu\text{m}$ column (cat no. 1893471). The analytical column output was coupled to a 5 cm × 20 $\mu\text{m}$ ID Sharp Singularity Fossil Ion Tech tapered emitter mounted in a custom microspray source heated to 45°C.

For each sample, peptides were eluted at 0.8 $\mu\text{L}/\text{min}$ using a 60-minute acetonitrile gradient as previously described.<sup>5</sup> Buffer A consisted of 0.1% formic acid in water, and buffer B consisted of 0.1% formic acid, 80% acetonitrile, and 20% water. Separate system suitability injections<sup>6</sup> were performed using a peptide mixture containing Pierce PRTCs and bovine serum albumin in 0.1% TFA.

DIA data were acquired on a Thermo Fisher Scientific Orbitrap Eclipse Tribrid mass spectrometer using 12 $m/z$ staggered precursor isolation windows.<sup>7,8</sup> The Orbitrap resolving power for all scans was 30,000 at $m/z$ 200. Each cycle consisted of one MS1 scan of $m/z$ 395-1,005 followed by 50 MS/MS scans. Automatic gain control was 100% for MS1 scans and 800% for MS/MS scans. MS/MS scans were acquired using a normalized collision energy of 27, assuming charge state 3. The staggered DIA method used two cycles: cycle 1 spanned $m/z$ 406.4347-994.7021, and cycle 2 spanned $m/z$ 400.4319-1000.7048. Spectra were acquired in centroid mode.

#### 3. Mass Spectrometry Data Processing

Raw mass spectrometry data were demultiplexed<sup>7,8</sup> and converted to mzML format using ProteoWizard’s msConvert<sup>9</sup> version 3.0.25168 with peak picking and demultiplexing enabled. An automated Nextflow workflow (nf-carafe-ai-ms version 85ee6c5 (https://nf-carafe-ai-ms.readthedocs.io/en/latest/)) was used to build an instrument-specific spectral library using the UniProt reviewed human proteome (UP000005640) plus common contaminants (https://www.thegpm.org/crap/), yeast enolase (P00924) and a single randomly picked liver sample spectral data file (012\_Ecl\_2025\_05\_28\_HL-135\_A5.mzML) as training data. This workflow implemented DIA-NN<sup>10</sup> version 1.8.1 and Carafe version 0.0.1<sup>11</sup>.

A second Nextflow workflow (nf-skyline-dia-ms, diann2-multiplate branch, git commit hash: 9ee1991e7534a59d4268760551dfb11bb2aed3ec available at https://github.com/mriddle/nf-skyline-dia-ms) was used to search the DIA data from the individual liver samples. This workflow used DIA-NN version 2.1.0 for searching along with the Carafe-generated library from the previous step. DIA-NN results were imported into Skyline as part of the workflow, where peak boundaries were automatically refined using Skyline’s “Impute boundaries for missing peaks” function.<sup>12</sup> Protein grouping was performed in Skyline using IDPicker-based protein association logic, in which proteins matching the same set of peptides were grouped as indistinguishable protein groups as previously described.<sup>5</sup> Skyline was set to remove peptides that mapped to more than one protein after which peptide and protein level normalization was performed using a third Nextflow workflow (https://github.com/uw-maccosslab/nf-dia-batch-correction). This workflow applied median normalization at the precursor (peptide) and protein levels and output .tsv files containing $\log_2(\text{area}+1)$ peak area values where area is the sum of the total area of all MS/MS transitions for a given peptide (peptide level) or all measured peptides in each protein (protein level). Normalized data from this pipeline were used in all downstream analyses. An otherwise identical Skyline document in which peptides mapping to more than one protein were retained (“keep\_dups”) was also generated as an additional resource for users interested in proteins for which informative peptides may be shared among multiple proteins; all analyses presented here used the “throw\_dups” dataset.

Skyline files, pipeline configuration files, and output from all pipelines are available on Panorama Public<sup>13</sup> using the URL: https://panoramaweb.org/uw-human-liver-bank-s9.url) and were assigned the ProteomeXchange<sup>14</sup> ID: PXD085173. Yeast enolase peptides, PRTC peptides, and pooled liver reference samples were used to evaluate sample processing, LC-MS/MS performance, and run-order effects. The pooled liver reference samples, generated from all 61 individual liver samples and included once per row in the 96-well KingFisher digest plate, were used as within-plate reference samples to assess technical reproducibility and potential row- or run-order-associated variation.

#### VIII. Development of minimal physiologically based pharmacokinetic model for prediction of metabolite formation clearance

As the metabolism only happens in the intracellular water space, the following ordinary differential equations were integrated into the previously established liver model<sup>15</sup> without clearance term in the central compartment.

Liver model without FABP1-CYP interaction (no $K_i$) was built using equations 1-7 and 16-21:

$$\frac{d[\text{Drug}]}{dt} = -T_1 - T_3 \quad (1)$$

$$\frac{d[\text{CYP}]}{dt} = -T_1 + T_2 - T_5 + T_6 \quad (2)$$

$$\frac{d[\text{FABP1}]}{dt} = -T_3 - T_4 + T_6 \quad (3)$$

$$\frac{d[\text{Drug} - \text{CYP}]}{dt} = T_1 - T_2 - T_4 \quad (4)$$

$$\frac{d[\text{Drug} - \text{FABP1}]}{dt} = T_3 - T_5 \quad (5)$$

$$\frac{d[\text{Drug} - \text{FABP1} - \text{CYP}]}{dt} = T_4 + T_5 - T_6 \quad (6)$$

$$\frac{d[\text{Metabolite}]}{dt} = T_2 + T_6 \quad (7)$$

Liver model with FABP1-CYP interaction (with $K_i$) was built using equations 8-15 and 16-22:

$$\frac{d[\text{Drug}]}{dt} = -T_1 - T_3 \quad (8)$$

$$\frac{d[\text{CYP}]}{dt} = -T_1 + T_2 - T_5 - T_7 \quad (9)$$

$$\frac{d[\text{FABP1}]}{dt} = -T_3 - T_4 - T_7 \quad (10)$$

$$\frac{d[\text{Drug} - \text{CYP}]}{dt} = T_1 - T_2 - T_4 \quad (11)$$

$$\frac{d[\text{Drug} - \text{FABP1}]}{dt} = T_3 - T_5 \quad (12)$$

$$\frac{d[\text{Drug} - \text{FABP1} - \text{CYP}]}{dt} = T_4 + T_5 - T_6 \quad (13)$$

$$\frac{d[\text{FABP1} - \text{CYP}]}{dt} = T_6 + T_7 \quad (14)$$

$$\frac{d[\text{Metabolite}]}{dt} = T_2 + T_6 \quad (15)$$

The T terms with subscripts were defined as follows:

$$T_1 = k_{\text{on},\text{dp}} \cdot [\text{Drug}] \cdot [\text{CYP}] - k_{\text{off},\text{dp}} \cdot [\text{Drug} - \text{CYP}] \quad (16)$$

$$T_2 = k_{\text{cat}} \cdot [\text{Drug} - \text{CYP}] \quad (17)$$

$$T_3 = k_{\text{on},\text{df}} \cdot [\text{Drug}] \cdot [\text{FABP1}] - k_{\text{off},\text{df}} \cdot [\text{Drug} - \text{FABP1}] \quad (18)$$

$$T_4 = k_{\text{on},\text{dpf}} \cdot [\text{Drug} - \text{CYP}] \cdot [\text{FABP1}] - k_{\text{off},\text{dpf}} \cdot [\text{Drug} - \text{FABP1} - \text{CYP}] \quad (19)$$

$$T_5 = k_{\text{on},\text{dfp}} \cdot [\text{Drug} - \text{FABP1}] \cdot [\text{CYP}] - k_{\text{off},\text{dfp}} \cdot [\text{Drug} - \text{FABP1} - \text{CYP}] \quad (20)$$

$$T_6 = \beta k_{\text{cat}} \cdot [\text{Drug} - \text{FABP1} - \text{CYP}] \quad (21)$$

$$T_7 = k_{\text{on},\text{pp}} \cdot [\text{CYP}] \cdot [\text{FABP1}] - k_{\text{off},\text{pp}} \cdot [\text{FABP1} - \text{CYP}] \quad (22)$$

The $k_{\text{on}}$ and $k_{\text{off}}$ terms were defined as follows in the model:

$$K_m = \frac{k_{\text{off},\text{dp}}}{k_{\text{on},\text{dp}}} \quad (23)$$

$$K_d = \frac{k_{\text{off},\text{df}}}{k_{\text{on},\text{df}}} \quad (24)$$

$$\alpha K_m = \frac{k_{\text{off},\text{dfp}}}{k_{\text{on},\text{dfp}}} \quad (25)$$

$$\alpha K_d = \frac{k_{\text{off},\text{dpf}}}{k_{\text{on},\text{dpf}}} \quad (26)$$

$$K_i = \frac{k_{\text{off},\text{pp}}}{k_{\text{on},\text{pp}}} \quad (27)$$

In equations 1-27, [Drug] is the concentration of free drug, [CYP] is the concentration of free CYP2C9, [FABP1] is the concentration of free FABP1, [Drug-CYP] is the concentration of drug bound in the CYP2C9 active site, [Drug-FABP1] is the concentration of the drug bound to FABP1, [Drug-FABP1-CYP] is concentration of the ternary complex, [FABP1-CYP] is the concentration of the protein-protein complex, and [Metabolite] is the concentration of metabolites formed. Each $T_i$ term represents the net flux in concentration per time. $T_1$ is the net binding of free drug to free

CYP2C9, $T_2$ is the catalytic turnover of the Drug-CYP2C9 complex to metabolite, $T_3$ is the net binding of free drug to free FABP1, $T_4$ is the net binding of FABP1 to the Drug-CYP2C9 complex, $T_5$ is the net binding of CYP2C9 to Drug-FABP1 complex, $T_6$ is the catalytic turnover of the ternary complex to metabolite, and $T_7$ is the net binding of free CYP2C9 to free FABP1. $k_{on,dp}$ and $k_{off,dp}$ are the association and dissociation rate constants of drug with CYP2C9, $k_{on,df}$ and $k_{off,df}$ are the association and dissociation rate constants of drug with FABP1, $k_{on,dpf}$ and $k_{off,dpf}$ are the association and dissociation rate constants of FABP1 with Drug-CYP2C9 complex, $k_{on,dfp}$ and $k_{off,dfp}$ are the association and dissociation rate constants of CYP2C9 with Drug-FABP1 complex, and $k_{on,pp}$ and $k_{off,pp}$ are the association and dissociation rate constants of CYP2C9 with FABP1. $k_{cat}$ is the turnover number of the binary Drug-CYP2C9 complex. $\alpha$ and $\beta$ are the dimensionless parameters estimated in **Figure 5**. $K_m$ is the dissociation equilibrium constant of drug from CYP2C9 and assumed to be the same as Michaelis constant under rapid equilibrium. $K_d$ is the dissociation equilibrium constant of drug from FABP1. $\alpha K_d$ is the dissociation equilibrium constant of FABP1 from the ternary complex. $\alpha K_m$ is the dissociation equilibrium constant of CYP2C9 from the ternary complex. $K_i$ is the dissociation equilibrium constant of the CYP2C9-FABP1 complex.

All simulations were run to steady state and hence are insensitive to $k_{on}$ and $k_{off}$ values. The experimentally determined $k_{on}$ in Supplementary Table 2 for DAUDA-FABP1 binding is close to diffusional rate limited, so drug-protein and protein-protein association process were assumed to be diffusional rate limited ($360,000 \mu M^{-1} hr^{-1}$).<sup>16</sup>

The developed liver models can be found at GitHub: Isoherranen-Lab.

**Supplementary Table 6. Physiological parameters used in the developed liver distribution models.** Physiological parameters are based on a 70-kilogram healthy adult human. Table retrieved from Wen and Isoherranen (2026) *JPET*.<sup>15</sup>

|  | Human |
| --- | --- |
| ***Liver Volume (L)***<sup>17</sup> | 1.8 |
| Liver blood<sup>17</sup> | 0.2 |
| Liver tissue<sup>17</sup> | 1.6 |
| Liver – space of disse<sup>17,18</sup> | 0.26 |
| Liver – intracellular space<sup>17,18</sup> | 0.92 |
| ***Fraction of Liver Wet Weight*** |  |
| Water<sup>18</sup> | 0.73 |
| Neutral lipids (f<sub>NL</sub>)<sup>18</sup> | 0.0348 |
| Neutral phospholipids (f<sub>NP</sub>)<sup>18</sup> | 0.0252 |
| Extracellular water (f<sub>ECW</sub>)<sup>18</sup> | 0.161 |
| Intracellular water (f<sub>ICW</sub>)<sup>18</sup> | 0.573 |
| ***Fraction of Plasma Wet Weight*** |  |
| Neutral lipids (f<sub>p,NL</sub>)<sup>18</sup> | 0.0032 |
| Neutral phospholipids (f<sub>p,NP</sub>)<sup>18</sup> | 0.0021 |
| ***pH*** |  |
| Plasma<sup>19</sup> | 7.4 |
| Extracellular water<sup>19</sup> | 7.4 |
| Intracellular water<sup>19</sup> | 7.2 |
| ***Albumin Ratio*** |  |
| Extracellular water-to-plasma<sup>20,21</sup> | 0.63 |
| Liver-to-plasma<sup>18,21</sup> | 0.101<sup>a</sup> |
| ***Albumin Concentration (μM)*** |  |
| Plasma<sup>22</sup> | 640 |
| Liver – extracellular water<sup>20–22</sup> | 403<sup>b</sup> |

<sup>a</sup>Liver-to-plasma albumin ratio is calculated as $f_{ECW} \times$ extracellular-to-plasma albumin ratio.

<sup>b</sup>Albumin concentration in the extracellular water space in liver is calculated as the plasma albumin concentration × extracellular water-to-plasma albumin ratio.

**Supplementary Table 7. Model input parameters for prediction of *in vivo* 4'OH-diclofenac and 4'OH-tolbutamide clearance.**

|  | 4'OH-diclofenac | 4'OH-tolbutamide |
| --- | --- | --- |
| CYP2C9 ($\mu\text{M}$)<sup>a</sup> | 9 | 9 |
| $k_{\text{cat}}$ ($\text{hr}^{-1}$) | 390 | 130 |
| $K_{\text{m,u}}$ ($\mu\text{M}$) | 6.6 | 664.6 |
| $K_{\text{d}}$ ($\mu\text{M}$) | 5.2 | 7.4 |
| ***Without apo-FABP1-CYP2C9 interaction*** |  |  |
| $\alpha$ | 2.4 | 0.4 |
| $\beta$ | 1.6 | 10 |
| ***With apo-FABP1-CYP2C9 direct interaction*** |  |  |
| $\alpha$ | 0.03 | 0.01 |
| $\beta$ | 1.1 | 2.1 |
| $K_{\text{i}}$ ($\mu\text{M}$) | 0.5 | 0.2 |

<sup>a</sup>Intracellular CYP2C9 concentration was calculated to be 9 $\mu\text{M}$ taking the CYP2C9 expression level of 127 pmol/mg microsomal protein, an average microsomal protein yield of 40 mg/g liver, and fraction of liver wet weight as intracellular water into account.<sup>15,18,23,24</sup>
